# GFRAL neurons remodel intestinal structure and type 2 immunity

**DOI:** 10.64898/2026.10.28.762646

**Authors:** Daniëlle van de Lisdonk, Onur Eskiocak, Zachary Kerner, Wuqiang Guan, Semir Beyaz, Helmut W. Kessels, Bo Li

**Affiliations:** Cold Spring Harbor Laboratory; Cold Spring Harbor, NY 11724, USA; Swammerdam Institute for Life Sciences, Amsterdam Neuroscience, University of Amsterdam; Amsterdam, the Netherlands; Graduate Program in Genetics, Stony Brook University; Stony Brook, NY, USA; Laboratory of Mucosal Immunology, Rockefeller University; New York, NY, USA; Westlake Laboratory of Life Sciences and Biomedicine; Hangzhou, 310024 Zhejiang, China; School of Life Sciences, Westlake University; Hangzhou, 310024 Zhejiang, China; Institute of Biology, Westlake Institute for Advanced Study; Hangzhou, 310024 Zhejiang, China

## Abstract

The brain has emerged as a key regulator of immune functions, but the underlying neural mechanisms remain poorly understood. Here we show that brainstem GFRAL-expressing neurons prime type 2 immunity and shape intestinal epithelial structure in mice. Type 2 inflammation results in activation of GFRAL neurons and potentiation of their excitatory synaptic inputs. Enhancing GFRAL neuron excitability is sufficient to remodel intestinal type 2 immunity through a polysynaptic vagal brain-gut circuit, expanding tuft and goblet cell populations and increasing intestinal type 2 cytokine expression; while inhibiting GFRAL neurons leads to largely opposite changes. Accordingly, manipulating GFRAL neuron function reshapes intestinal structure and pathophysiology in food allergy, helminth infection and colitis models. Thus, GFRAL neurons orchestrate type 2 immune responses to facilitate recovery from intestinal diseases.

## Main Text

The intestine is an organ that is continuously exposed to dietary antigens, commensal microbes and pathogens, and where immune responses must balance host defense with preservation of epithelial integrity (*1–5*). Type 2 immunity is central to this balance, driving responses to parasites (e.g., helminths), food allergens and epithelial injury through activation of group 2 innate lymphoid cells (ILC2s), expansion of tuft and goblet cells, and production of cytokines such as interleukin (IL)-4, IL-5 and IL-13 (*6–10*). Together, these pathways promote helminth clearance, mucus production and tissue repair, while their dysregulation contributes to diseases like allergic diseases, barrier dysfunctions and inflammatory bowel disease (IBD) (*6, 7, 11*).

In parallel to its immune complexity, the intestine is innervated by the nervous system, including extrinsic sympathetic and parasympathetic inputs and the intrinsic enteric nervous system, which regulate intestinal nutrient sensing, motility and secretion (*3–5*). More recent work indicates that the central nervous system also monitors and modulates immune function and epithelial remodeling in the gut (*11–14*). By integrating inflammatory, epithelial and immune derived signals and providing feedback, the nervous system plays a critical role in maintaining intestinal homeostasis (*15–18*). When this interaction is disrupted, intestinal inflammation is exacerbated and disease outcomes are worsened (*19, 20*). Nevertheless, how the brain monitors immune state and translates the state signals into protective intestinal responses remains largely undefined.

Growth differentiation factor 15 (GDF15) is a stress-induced cytokine that is elevated in a range of pathological conditions, including infection, inflammation, cancer and metabolic diseases (*21–25*). Particularly, increased circulating GDF15 has been reported in mouse models of food allergy and other type 2-biased immune states (*26–28*). GDF15 signals through its only known receptor, GDNF family receptor alpha-like (GFRAL), which is expressed exclusively in a restricted population of neurons in the area postrema (AP) and the adjacent nucleus of the solitary tract (NTS) in the brainstem (*29–32*). Interestingly, the AP is a circumventricular organ that lacks the conventional blood-brain barrier (BBB), permitting circulating factors and cytokines to directly access its neurons, including GFRAL-expressing (GFRAL^+^) neurons (*33–35*).These findings suggest that GFRAL^+^ neurons in the AP may function as sensors of systemic stress signals. In the present study, we tested whether these neurons might function not only as sensors, but also as effectors capable of modulating peripheral physiology, such as intestinal immunity.

### Type 2 immunity promotes GFRAL^+^ neuron activity

Previous studies have shown that food allergy in mouse models leads to elevated plasma GDF15 levels (*26–28*). Because GFRAL is the only known receptor for GDF15 (*29–32*), we asked whether food allergy results in activation of GFRAL^+^ neurons. We first established the food allergy model in C57BL/6 mice, whereby mice were sensitized by repeated subcutaneous (s.c.) administration of ovalbumin (OVA) mixed with the adjuvant aluminum (alum) hydroxide (Fig. 1A; Methods). Control mice received phosphate-buffered saline (PBS) and alum. Both groups of mice were subsequently challenged with OVA via oral gavage. At three hours following the final OVA challenge, we examined the expression of Fos — an immediate early gene product that is an indicator of neuronal activation (*36, 37*) — with immunohistochemistry in brain sections. We observed that, in the sensitized mice relative to the control mice, there was an increase in Fos^+^ cells in the AP and NTS, and an increase in Fos^+^ cells among GFRAL^+^ neurons (Fig. 1B, C). In contrast, the parabrachial nucleus (PBN) and central amygdala (CeA) showed little Fos induction at this time point (fig. S1A, B). This difference is consistent with prior work showing that the PBN and CeA are engaged during the rapid vagal-mediated aversive responses in food allergy (*26, 38, 39*), while AP activity can reflect slower, blood-borne cytokine signaling (*40*).

**Fig. 1.**
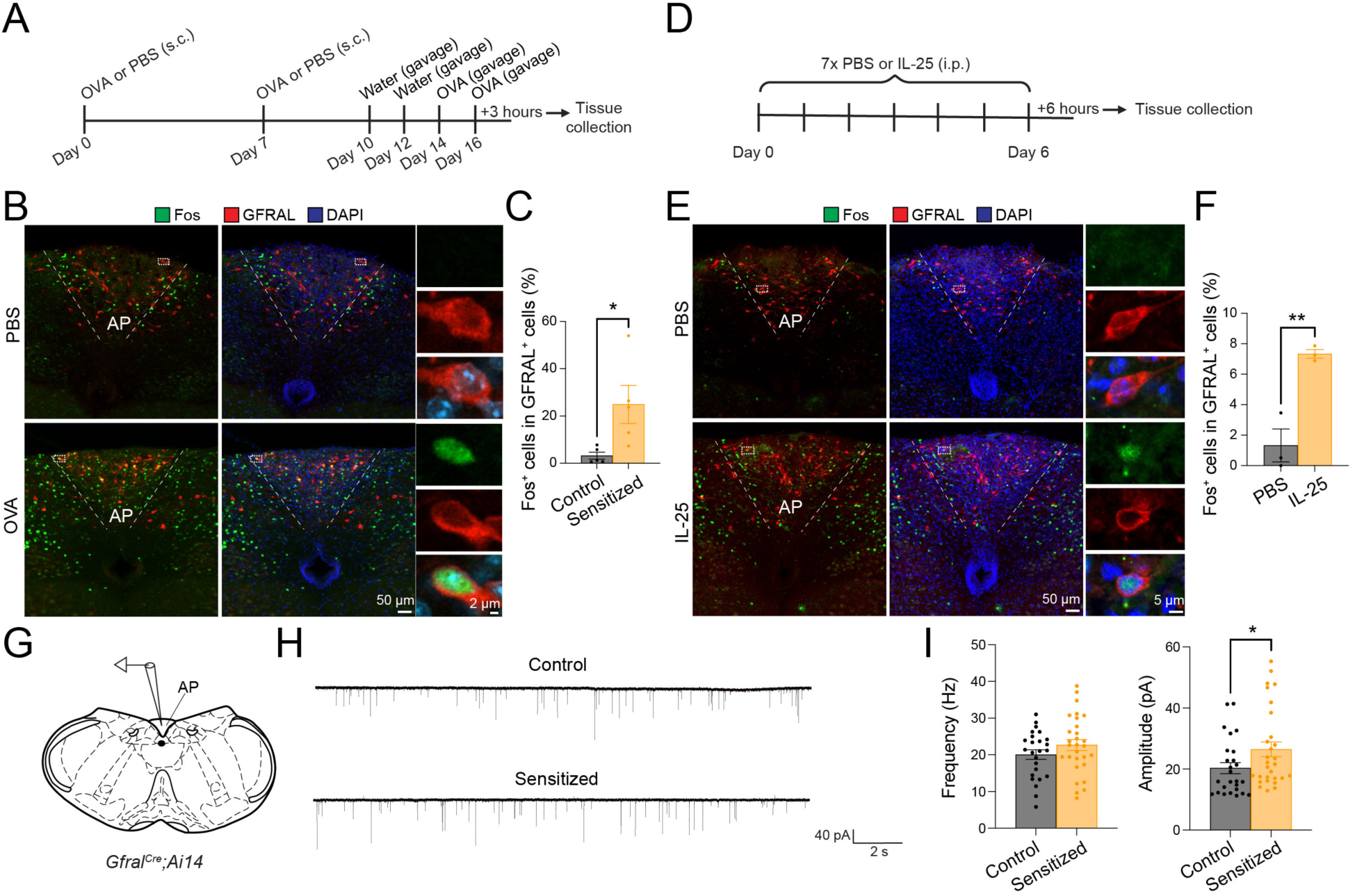
Type 2 immune responses activate GFRAL neurons. **(A)** Experimental timeline for allergic sensitization in C57BL/6 mice treated with either OVA + alum or PBS + alum (control), followed by two days of water gavage and two subsequent OVA gavages. **(B)** Immunohistochemical images of the AP stained for Fos and GFRAL after OVA gavage; scale bar, 50 and 2 μm. **(C)** Quantification of Fos+ GFRAL neurons (PBS + alum, n = 5, OVA + alum, n = 5; *P = 0.0295; Unpaired t test). **(D)** Experimental timeline for IL-25 or PBS treatment in C57BL/6 mice for seven consecutive days. **(E)** Immunohistochemical images of the AP stained for Fos and GFRAL after IL-25 or PBS treatment; scale bar, 50 and 5 μm. **(F)** Quantification of Fos+ GFRAL neurons (PBS, n = 3, IL25, n = 3; **P = 0.0057; Unpaired t test). **(G)** Diagram showing the AP in a coronal brain section for electrophysiological recording. **(H)** Representative miniature EPSC traces from GFRAL neurons in PBS + Alum mice (top) and OVA + Alum mice (bottom). **(I)** Quantification of miniature EPSC frequency (left), amplitude (right) (PBS + alum, n = 29 cells / 5 mice, OVA + alum, n = 37 cells / 6 mice; frequency, P = 0.7151 unpaired t test; amplitude, *P = 0.0381, Mann-Whitney test). Data are presented as mean s.e.m.

To determine whether GFRAL^+^ neurons are activated generally by type 2 immune stimuli, we next adopted a T helper 2 (Th2) cell-biased inflammation model induced by IL-25 injections (Fig. 1D; Methods). It is known that IL-25 plays an important role in coordinating type 2 immunity, increasing Th2 cytokine (IL-4, IL-5 and IL-13) production and inducing intestinal tuft and goblet cell expansion (*41–43*), processes critical for fighting against helminth infection (*44*). Notably, IL-4 and IL-13 have been shown to stimulate GDF15 production via STAT6- dependent signaling in adipose tissue, raising the possibility that Th2 inflammation may engage GDF15-GFRAL signaling pathways (*45*). We assessed Fos expression six hours following the final IL-25 injection. Consistent with the above result, IL-25 increased Fos^+^ cells among GFRAL^+^ neurons (Fig. 1E, F) but did not increase Fos^+^ cells in the PBN or CeA (fig. S1C, D). Therefore, both antigen and cytokine type-2-immune stimuli activate GFRAL^+^ neurons, supporting the idea that these neurons function as a CNS sensor of systemic type 2 inflammatory states.

We next assessed whether type 2 immune stimuli can cause functional changes in GFRAL^+^ neurons. To this end, we used *Gfral^Cre^;Ai14* mice — in which GFRAL^+^ neurons express the red fluorescence protein tdTomato — for the food allergy model described above, and prepared acute brain slices from these mice after the last challenge with OVA (Fig. 1G; Methods). We recorded miniature excitatory postsynaptic currents (mEPSCs) from GFRAL^+^ neurons, which were identified based on tdTomato fluorescence. Notably, the amplitude of the mEPSCs in OVA group was larger than that in control group (Fig. 1H, I). This result shows that food allergy is accompanied by enhanced excitatory synaptic transmission onto GFRAL^+^ neurons, which likely contributes to their heightened activity.

### Endogenous GFRAL^+^ neuron activity facilitates intestinal type 2 immunity

We wondered whether suppressing the activity of GFRAL^+^ neurons would reduce food allergy. In type 2 immunity, intestinal tuft cells sense allergens (e.g., those originated from foods or parasites) and release IL-25 and leukotrienes that stimulate nearby ILC2s (*6, 8–10, 43*). Activated ILC2s produce IL-4, IL-5 and IL-13, which drive stem cells to expand tuft and goblet cell populations (*6*). The resulting increase in goblet cells boosts mucus production, helping to defend the intestine against parasitic infection.

To selectively inhibit GFRAL^+^ neurons, we infected them with an adeno-associated virus (AAV) expressing the tetanus toxin light chain (TeLC), which blocks neurotransmitter release, or GFP as the control (Fig. 2A). These mice were subjected to the food allergy paradigm as described above. Remarkably, TeLC normalized the allergy-associated increase in plasma IgE levels (Fig. 2B). This manipulation did not reduce GDF15 levels under this allergic condition; rather, it increased plasma and intestinal GDF15 levels (Fig. 2C). No differences in IL-4, IL-5 or IL-13 mRNA levels were observed in the small intestine of different groups (fig. S2A). These results suggest that endogenous GFRAL^+^ neuron activity facilitates IgE-mediated responses during food allergy.

**Fig. 2.**
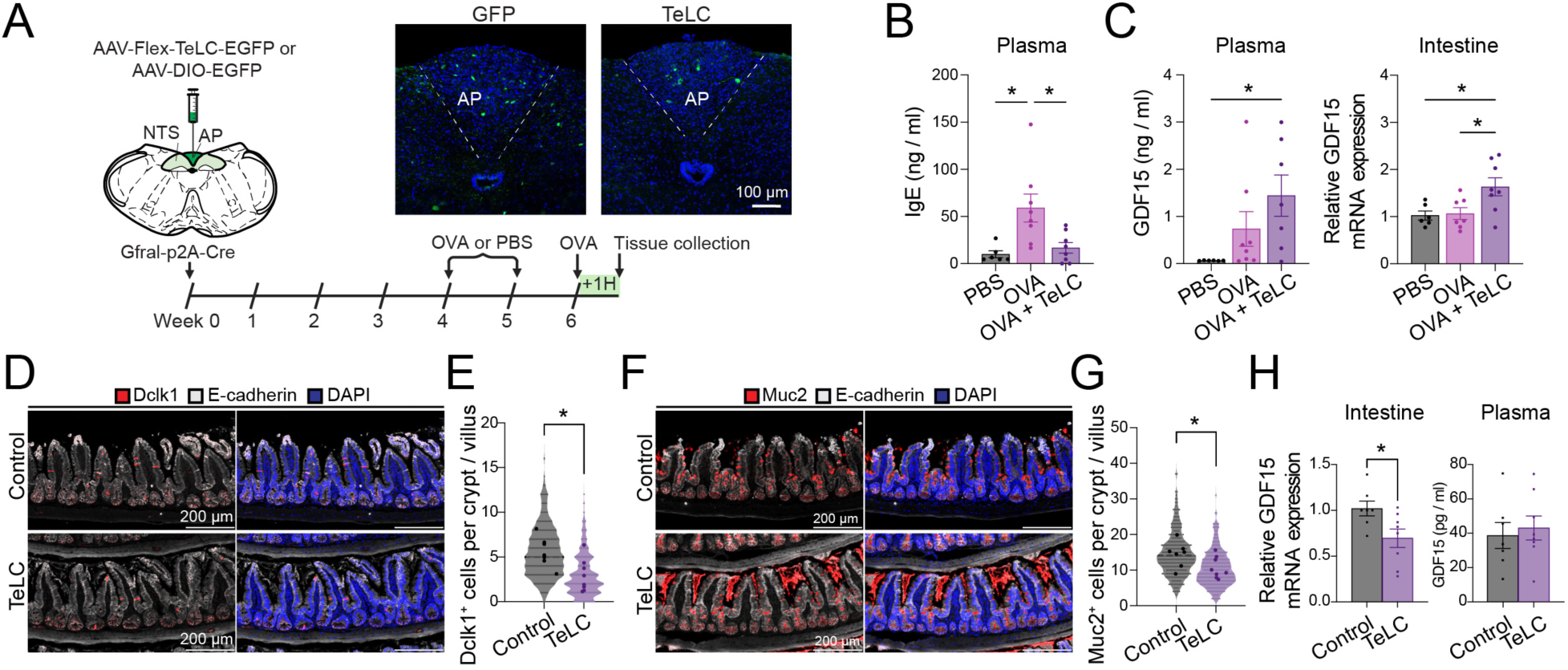
Inhibition of GFRAL neurons decreases allergen-specific antibody production. **(A)** Timeline of the experiments, in which TeLC or eGFP was injected at week 0. Mice were treated with either OVA + alum or PBS + alum (control) starting at week 4, followed by two days of water gavage and one subsequent OVA gavage. Example immunohistochemical images of the AP showing expression of eGFP or TeLC; scale bar, 100 μm. **(B)** Plasma IgE concentrations in PBS + alum mice (n = 6), OVA + alum mice (n = 8), and TeLC + OVA + alum mice (n = 8) (PBS+alum vs OVA+alum, *P = 0.0113; PBS+alum vs OVA+alum+TeLC, P = 0.9578; OVA+alum vs OVA+alum+TeLC, *P = 0.0188; one-way ANOVA followed by Sidak’s post hoc test). **(C)** (Left) Plasma GDF15 concentrations in control (n = 6), GFP with OVA + alum (n = 8), and TeLC with OVA + alum treated mice (n = 7) (PBS+alum vs OVA+alum, P = 0.4854; PBS+alum vs OVA+alum+TeLC, *P = 0.0465; OVA+alum vs OVA+alum+TeLC, P = 0.4027; one-way ANOVA followed by Sidak’s post hoc test). (Right) Expression of GDF15 mRNA in the small intestine of TeLC mice (PBS + alum (n=6) vs. OVA + alum (n=7), P = 0.9806; PBS + alum vs. OVA + alum + TeLC (n = 8), *P = 0.0310; OVA + alum vs. OVA + alum + TeLC, *P = 0.0366). **(D)** Immunohistochemical images of the ileum in TeLC mice, showing the expression of Dclk1 and E-cadherin; scale bar, 200 μm. **(E)** Quantification of Dclk1+ cells per crypt/villus structure (control, n = 7, TeLC n = 8; *P = 0.0108; Unpaired t test). **(F)** Immunohistochemical images of the ileum in TeLC mice, showing the expression of Muc2 and E-cadherin; scale bar, 200 μm. **(G)** Quantification of Muc2+ cells per crypt/villus structure (control, n = 7, TeLC n = 8; *P = 0.0473; Unpaired t test). **(E, G)** Each small point represents the number of Dclk1+ or Muc2+ cells per crypt/villus, and each larger point indicates the average value for an individual mouse. Statistical analyses were performed on the per-mouse averages. **(H)** Left, expression of GDF15 mRNA in the small intestine of TeLC mice (control, n = 7, TeLC n = 8; *P = 0.0282; Unpaired t test). Right, plasma GDF15 concentrations in control and TeLC mice (control, n = 7, TeLC, n = 8; P = 0.6740; Unpaired t test). Data are presented as mean s.e.m.

We also assessed metabolic parameters in these mice, including intestinal and colonic length, as well as skeletal muscle, inguinal white adipose tissue (iWAT) and brown adipose tissue (BAT) mass, which were largely similar across groups, except that the TeLC mice exhibited increased BAT mass (fig. S2B-D). This latter observation suggests a role of endogenous GFRAL^+^ neuron activity in regulating BAT.

To determine whether GFRAL^+^ neurons influence type 2 immunity even under immune- naïve conditions, we prepared new cohorts of TeLC mice and GFP mice (fig. S3A). Notably, the number of cells expressing Dclk1 or Muc2, which are well-established markers for tuft cells and goblet cells, respectively, was markedly decreased in the small intestine of TeLC mice compared with GFP mice (Fig. 2D-G). Notably, TeLC mice showed decreased intestinal GDF15, although their plasma GDF15 remained unchanged (Fig. 2H). TeLC mice had normal levels of plasma immunoglobulin E (IgE) and intestinal type 2 cytokines (fig. S3B) and had normal body weight and intact ileal villus length and crypt length (fig. S3C-E). Further metabolic analysis confirmed that TeLC mice had normal food and water intake, respiratory exchange ratio (RER), energy expenditure (EE), and locomotion (fig. S3F-K). Thus, inhibiting GFRAL^+^ neurons selectively reduced intestinal tuft cells and goblet cells, as well as intestinal GDF15 levels, indicating that these neurons regulate type 2 immunity-related processes under baseline conditions. The results that inhibiting GFRAL^+^ neurons either increases or reduces intestinal GDF15 levels under challenged or immune naïve conditions, respectively, suggest that GFRAL^+^ neurons bidirectionally regulate intestinal GDF15 production in an immune state-dependent manner.

### GFRAL^+^ neurons remodel intestinal type 2 immune program

We next examined the effects of promoting GFRAL^+^ neuron activity on type 2 immunity. We chronically increased GFRAL^+^ neuron excitability, aimed at mimicking the chronic effects that might be imposed by type 2 immune conditions. Two strategies were used. The first one was based on the *Rosa26-stop^flox^-GluA2^mut^* mouse line, which conditionally expresses a mutant form of AMPA receptor GluA2 subunit (GluA2^mut^) (*46*). We bred *Gfral^Cre^;Rosa26-stop^flox^-GluA2^mut^*mice (referred to as “GluA2^mut^ mice”; Methods), in which GluA2^mut^ was selectively expressed in GFRAL^+^ neurons (fig. S4A-C). Their littermates without GluA2^mut^ expression were used as controls (referred to as “GluA2^control^ mice”). The GluA2^mut^ potentiated excitatory synaptic transmission onto GFRAL^+^ neurons (fig. S4D-G), consistent with previous results from other neuronal types (*46–49*). The second strategy was to use the bacterial voltage-gated sodium channel (NaChBac) (*50–54*) to increase the intrinsic excitability of GFRAL^+^ neurons. For this purpose, we injected the AP of *Gfral^Cre^* mice with an AAV expressing NaChBac (or GFP as a control) in a Cre-dependent manner (fig. S4H, I).

Notably, we found that the number of tuft cells was markedly increased in the small intestine of both the GluA2^mut^ mice and the NaChBac mice relative to their controls (Fig. 3A, B). Consistently, goblet cells were also increased in these mice (Fig. 3C, D). Moreover, female GluA2^mut^ mice, as well as male and female NaChBac mice, showed significantly increased intestinal IL-4 mRNA expression (Fig. 3E, G). Both models displayed elevated intestinal GDF15 mRNA levels (Fig. 3F, H), although their plasma GDF15 levels remained unchanged (fig. S4J). Thus, elevating the synaptic or intrinsic excitability of GFRAL^+^ neurons in the brainstem, with GluA2^mut^ and NaChBac, respectively, promotes a type 2-biased epithelial and cytokine program in the intestine, indicating that GFRAL^+^ neurons actively regulate intestinal type 2 immunity.

**Fig. 3.**
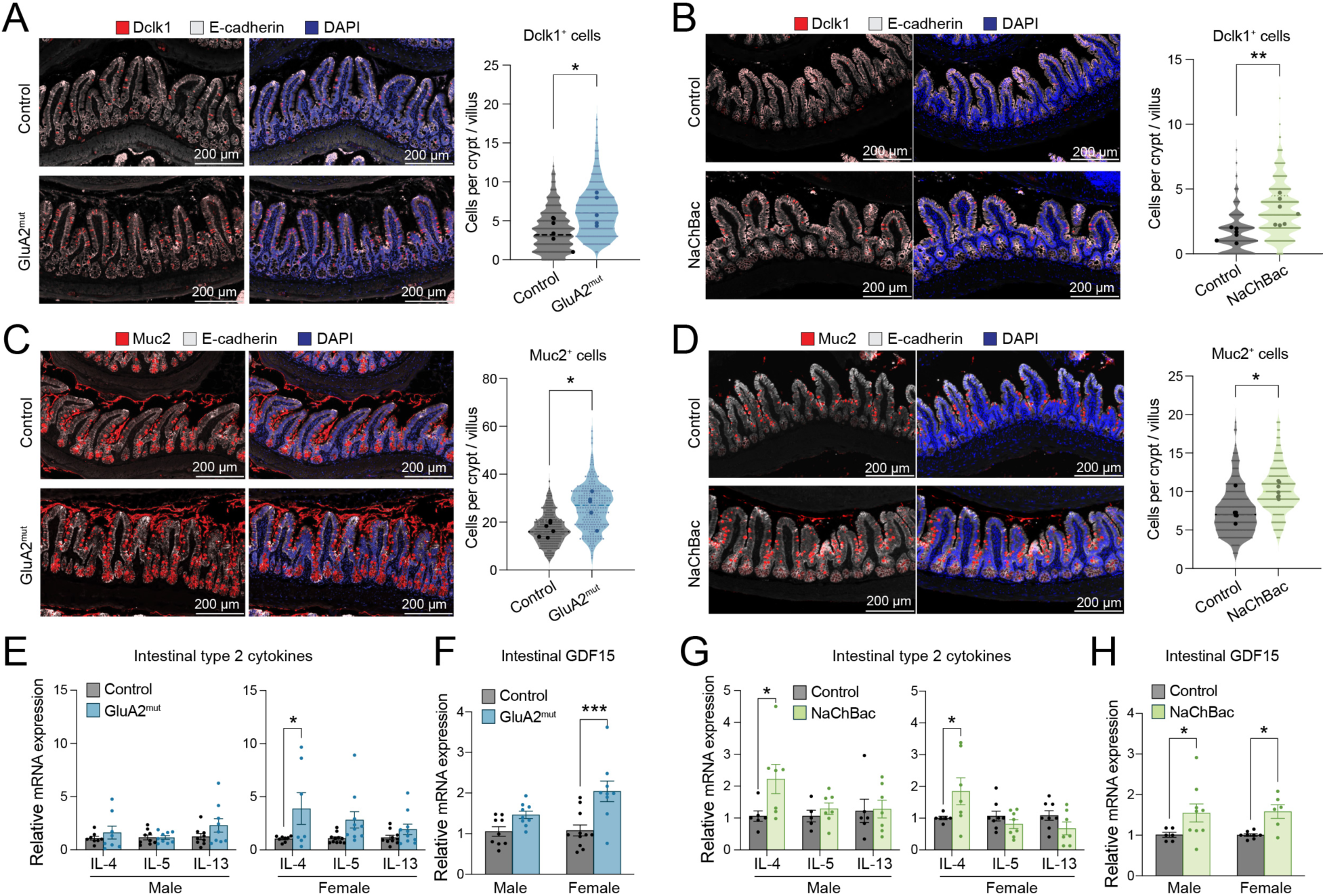
Activation of GFRAL neurons induces type 2 immune response in the intestine. **(A)** Immunohistochemical images of the ileum in GluA2^mut^ mice, showing the expression of Dclk1 and E-cadherin; scale bar, 200 μm. And quantification of Dclk1+ cells per crypt/villus structure. (control, n = 6, GluA2^mut^, n = 5; *P = 0.0460; Unpaired t test). **(B)** Immunohistochemical images of the ileum in NaChBac mice, showing the expression of Dclk1 and E-cadherin; scale bar, 200 μm. And quantification of Dclk1+ cells per crypt/villus structure. (control, n = 6, NaChBac, n = 7; **P = 0.0033; Unpaired t test). **(C)** Immunohistochemical images of the ileum in GluA2^mut^ mice, showing the expression of Muc2 and E-cadherin; scale bar, 200 μm. And quantification of Muc2+ cells per crypt/villus structure. (control, n = 6, GluA2^mut^, n = 5; *P = 0.0113; Unpaired t test). **(D)** Immunohistochemical images of the ileum in NaChBac mice, showing the expression of Muc2 and E-cadherin; scale bar, 200 μm. And quantification of Muc2+ cells per crypt/villus structure. (control, n = 6, NaChBac, n = 7; *P = 0.0221; Mann-Whitney test). **(A-D)** Each small point represents the number of Dclk1+ or Muc2+ cells per crypt/villus, and each larger point indicates the average value for an individual mouse. Statistical analyses were performed on the per-mouse averages. **(E)** Expression of IL-4, IL-5 and IL-13 mRNA in the small intestine of (left) male and (right) female GluA2^mut^ mice (male: IL-4 control, n = 8, GluA2^mut^, n = 8; P = 0.7625; IL-5 control, n = 9, GluA2^mut^, n = 9; P > 0.9999; IL-13 control, n = 9, GluA2^mut^, n = 9; P = 0.1814; Šídák’s multiple comparisons test) (female: IL-4 control, n = 8, GluA2^mut^, n = 7; *P = 0.0147; IL-5 control, n = 11, GluA2^mut^, n = 10; P = 0.1095; IL-13 control, n = 10, GluA2^mut^, n = 10; P = 0.7408; Šídák’s multiple comparisons test). **(F)** Expression of GDF15 mRNA in the small intestine of GluA2^mut^ male and female mice (male, control, n = 9, GluA2^mut^, n = 9; P = 0.1686; female, control, n = 11, GluA2^mut^, n = 9; ***P = 0.0002; Šídák’s multiple comparisons test). **(G)** Expression of IL-4, IL-5 and IL-13 mRNA in the small intestine of (left) male and (right) female NaChBac mice (male: IL-4 control, n = 6, NaChBac, n = 7; *P = 0.0320; IL-5 control, n = 5, NaChBac, n = 7; P = 0.9397; IL-13 control, n = 6, NaChBac, n = 7; P = 0.9988; Šídák’s multiple comparisons test) (female: IL-4 control, n = 6, NaChBac, n = 7; *P = 0.0188; IL-5 control, n = 8, NaChBac, n = 8; P = 0.7578; IL-13 control, n = 8, NaChBac, n = 7; P = 0.4430; Šídák’s multiple comparisons test). **(H)** Expression of GDF15 mRNA in the small intestine of NaChBac male and female mice (male, control, n = 6, NaChBac, n = 9; *P = 0.0500, female, control, n = 8, NaChBac, n = 6; *P = 0.0381; Šídák’s multiple comparisons test). Data are presented as mean s.e.m.

### GFRAL^+^ neurons regulate body weight independent of food intake

Previous studies show that acute activation of GFRAL^+^ neurons drives aversive responses and anorexia (*30, 55–59*), and that enhanced GFRAL^+^ neuron activity or GDF15/GFRAL signaling contributes to cancer cachexia in animal models (*40, 60–62*). Therefore, we also examined the GluA2^mut^ mice and NaChBac mice for metabolic phenotypes. Male GluA2^mut^ mice had reduced body weight accompanied by decreased iWAT and BAT mass compared with their controls (fig. S5A, B), although both male and female GluA2^mut^ mice had unaltered muscle mass and body length (fig. S5C, D), and normal body weight trajectories during a 4-week period (fig. S5E). Surprisingly, metabolic analyses (Methods) revealed normal food intake in GluA2^mut^ mice (Fig. 4A), suggesting that the reduced adiposity was due either to EE or nutrient absorption. Further analyses showed that the EE, RER, and locomotor activity in GluA2^mut^ mice remained normal (fig. 5F-H), indicating unaltered energy expenditure. However, these mice consumed less water and produced drier fecal pellets (Fig. 4B, C), though this alone was unlikely to explain the reduced fat mass. Notably, male GluA2^mut^ mice showed decreased triglyceride content in the small intestine (Fig. 4D), suggesting impaired nutrient absorption. Female GluA2^mut^ mice appeared unaffected (Fig. 4D), revealing a sex bias.

**Fig. 4.**
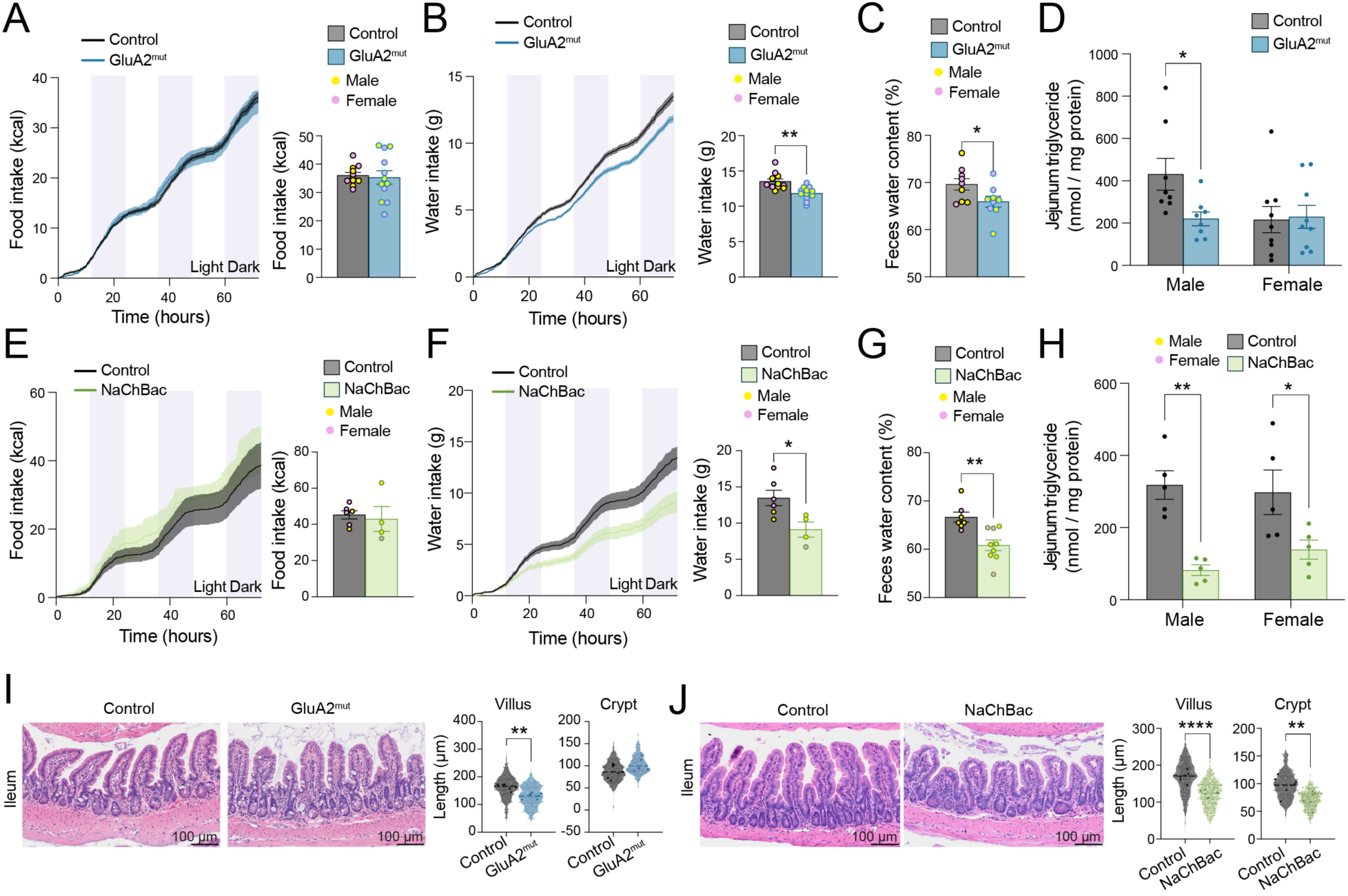
Activation of GFRAL neurons does not affect food intake but reduces nutrient absorption. **(A)** Food intake in GluA2^mut^ mice over time (Left) and average food intake (Right) (control, n = 11, GluA2^mut^, n = 12; P = 0.7770; Unpaired t test). **(B)** Water intake in GluA2^mut^ mice over time (Left) and average water intake (Right) (**P = 0.0011; Unpaired t test). **(C)** Fecal pellet water content in GluA2^mut^ mice (control, n = 9, GluA2^mut^, n = 9; *P = 0.0423; Unpaired t test). **(D)** Jejunal triglyceride levels in male and female GluA2^mut^ mice (Male, control, n = 8, GluA2^mut^, n = 8; *P = 0.0379, female, control, n = 9, GluA2^mut^, n = 9; P = 0.9843; Šídák’s multiple comparisons test). **(E)** Food intake in NaChBac mice over time (Left) and average food intake (Right) (P = 0.7173; Unpaired t test). **(F)** Water intake in NaChBac mice over time (Left) and average water intake (Right) (*P = 0.0242; Unpaired t test). **(G)** Fecal pellet water content in NaChBac mice (control, n = 7, NaChBac, n = 9; **P = 0.0021; Unpaired t test). **(H)** Jejunal triglyceride levels in male and female NaChBac mice (male control, n = 5, NaChBac, n = 5; female control, n = 5, NaChBac, n = 5) (Male, **P = 0.0013, female, *P = 0.0237; Šídák’s multiple comparisons test). **(A-H)** The last 72 hours were analyzed and plotted from the final week in the metabolic cages from GluA2^mut^ mice and NaChBac mice. **(I)** Representative HE images from Ileum in GluA2^mut^ mice; scale bar, 100 μm. And quantification of villus and crypt length. (control, n = 6, GluA2^mut^, n = 5; Villus, **P = 0.0043; Mann-Whitney test; Crypt, P = 0.1919; Unpaired t test). **(J)** Representative HE images from Ileum in NaChBac mice; scale bar, 100 μm. And quantification of villus and crypt length. (control, n = 6, NaChBac, n = 7; Villus ****P < 0.0001; Unpaired t test; Crypt, *P = 0.0083; Unpaired t test). **(I, J)** Each small point represents the number of Dclk1+ or Muc2+ cells per crypt/villus, and each larger point indicates the average value for an individual mouse. Statistical analyses were performed on the per-mouse averages. Data are presented as mean s.e.m.

**Fig. 5.**
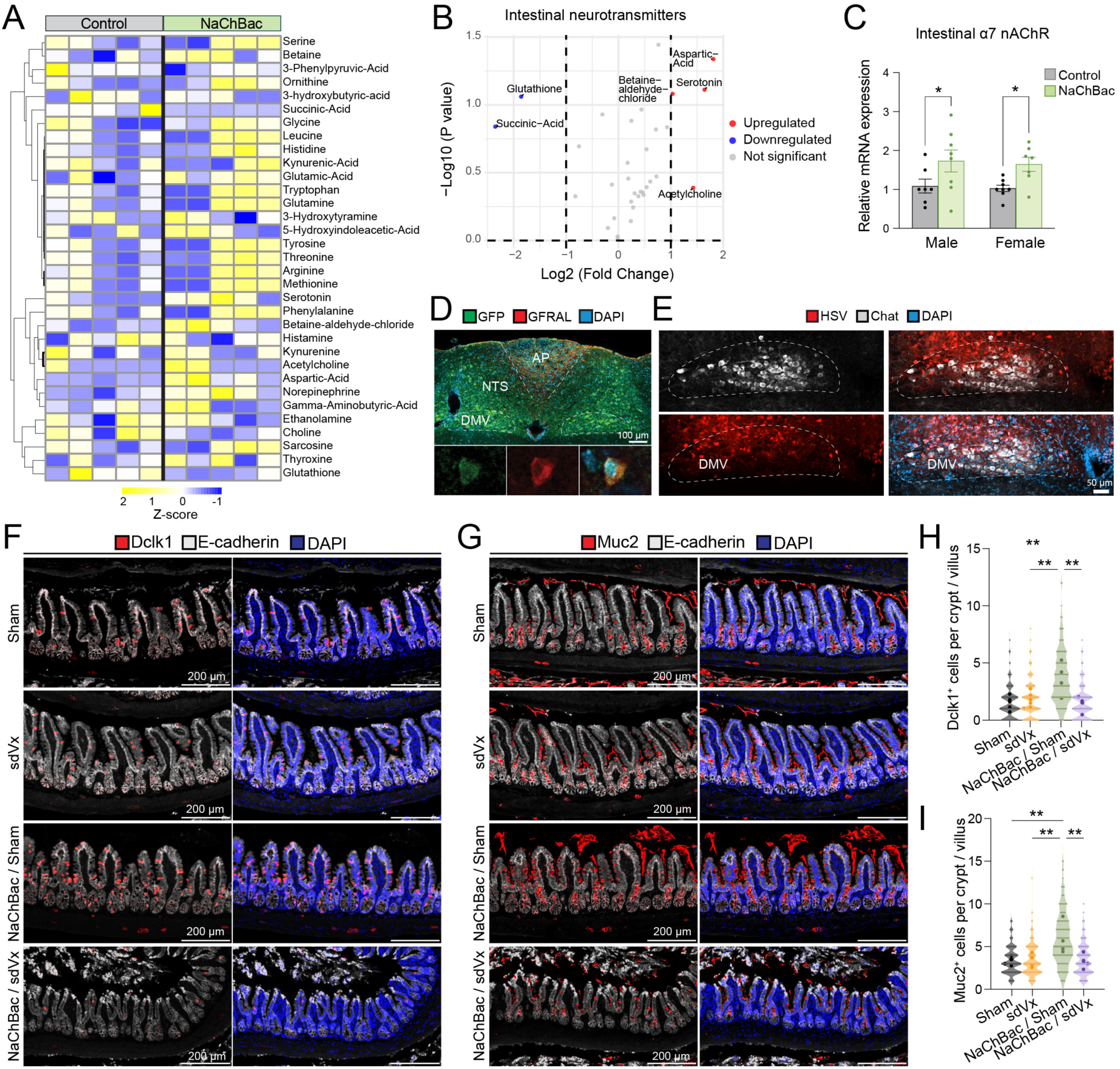
GFRAL neurons regulate intestinal type 2 immunity via the vagus nerve. **(A)** Heatmap showing the relative quantification of each metabolite in control and NaChBac mice in the small intestine, with yellow indicating higher levels and blue indicating lower levels. Cluster analysis was performed on the metabolites (control, n = 5, NaChBac, n = 5). **(B)** Volcano plot showing the relative differences and the statistical significance of metabolites between control and NaChBac mice in the small intestine. The X-axis represents the multiple change of the difference between the metabolites in different groups (log_2FoldChange), and the Y-axis represents the significance level of the difference (-log_10p-value). Blue and red dots denote up- and down-regulated metabolites. **(C)** Expression of α7nAChR mRNA in the small intestine of NaChBac male and female mice (Male, control, n = 7, NaChBac, n = 8; *P = 0.0312, Female, control, n = 8, NaChBac, n = 7; *P = 0.0262; Šídák’s multiple comparisons test). **(D)** Immunohistochemical images of the AP 5 days after PRV injections in the small intestine, stained for GFP and GFRAL; scale bars, 100 μm. **(E)** Immunohistochemical images of the DMV 3 days after HSV injections in the AP, stained for HSV and Chat; scale bars, 100 μm. **(F)** Immunohistochemical images of the ileum in Sham, sdVx, NaChBac+Sham and NaChBac+sdVx mice, showing the expression of Dclk1 and E-cadherin; scale bar, 200 μm. **(G)** Immunohistochemical images of the ileum in Sham, sdVx, NaChBac+Sham and NaChBac+sdVx mice, showing the expression of Muc2 and E-cadherin; scale bar, 200 μm. **(H)** Quantification of Dclk1+ cells per crypt/villus structure in sham (n = 6) vs sdVx (n = 6) vs NaChbac+sham (n = 4) vs NaChBac+sdVx (n = 6) (Sham vs. NaChBac + sham; **P = 0.0019, sdVx vs. NaChBac + sham **P = 0.0055, NaChBac + sham vs. NaChBac + sdVx; **P = 0.0021; Šídák’s multiple comparisons test). **(I)** Quantification of Muc2+ cells per crypt/villus structure in sham vs sdVx vs NaChbac+sham vs NaChBac+sdVx (Sham vs. NaChBac + sham; **P = 0.0047, sdVx vs. NaChBac + sham **P = 0.0041, NaChBac + sham vs. NaChBac + sdVx; **P = 0.0060; Šídák’s multiple comparisons test). **(H, I)** Each small point represents the number of Dclk1+ or Muc2+ cells per crypt/villus, and each larger point indicates the average value for an individual mouse. Statistical analyses were performed on the per-mouse averages. Data are presented as mean s.e.m.

Like GluA2^mut^ mice, the NaChBac mice had reduced iWAT (but in both sexes) and reduced BAT in males (fig. S6A). These mice also had normal food intake (Fig. 4E), locomotor activity and muscle mass (fig. S6B, C). Notably, like GluA2^mut^ mice, they consumed less water and had drier fecal pellets (Fig. 4F, G) and exhibited markedly reduced intestinal triglyceride levels (Fig. 4H). Unlike GluA2^mut^ mice, both male and female NaChBac mice exhibited progressive weight loss (fig. S6D) and decreased RER and EE during the dark phase (fig. S6E, F). These results show that enhancing GFRAL^+^ neuron activity with GluA2^mut^ or NaChBac can reduce body weight, adiposity and nutrient absorption without affecting food intake. These manipulations also reduce water intake, with NaChBac additionally reducing energy expenditure.

To understand how GFRAL^+^ neurons might influence nutrient absorption, we examined intestinal surface area, which is generally thought to correlate with fat absorption (*63, 64*). The GluA2^mut^ and NaChBac did not grossly affect intestine or colon length (fig. S7A-H). However, both models showed substantial reductions in ileal villus length, and NaChBac mice additionally displayed reduced ileal crypt depth (Fig. 4I, J). Moreover, although both models had normal villus length in the duodenum and jejunum, the NaChBac mice had markedly reduced crypt depth in these locations (fig. S7I-T). These findings suggest that heightened GFRAL^+^ neuronal activity impairs nutrient absorption in part by remodeling small intestinal morphology in ways that reduce absorptive surface area.

The manipulation with NaChBac in general resulted in stronger phenotypes than the manipulation with GluA2^mut^, likely because of the relative difference in chronicity of the two manipulations: whereas NaChBac was virally expressed during adult, GluA2^mut^ was expressed genetically and thus throughout development and adulthood. Developmental adaptations may compensate for the effects of GluA2^mut^ (also see below the Discussion section about the susceptibility of the GDF15/GFRAL system to desensitization).

### GFRAL^+^ neurons regulate type 2 immunity through a vagal brain-gut circuit

Inflammatory processes in the small intestine are influenced by inputs originating from the vagus nerve, the pelvic nerves, the gut’s sympathetic innervation, and the intrinsic neurons of the enteric nervous system (*3, 4, 15*). To identify the mechanisms through which GFRAL^+^ neurons shape type 2 immunity, we first asked whether GFRAL^+^ neuron activation would alter neurochemical signaling in the gut. Targeted neurotransmitter profiling of small intestinal tissue from NaChBac and control mice revealed a significant increase in acetylcholine (ACh) and serotonin in NaChBac mice (Fig. 5A, B). Interestingly, the expression of *Chrna7* (the gene encoding α7 nicotinic acetylcholine receptors (nAChRs)) was also increased in the small intestine of NaChBac mice (Fig. 5C). The vagus nerve releases ACh, which activates α7 nAChRs on immune cells to promote anti-inflammatory signaling (*65–69*). In addition, ACh released from the vagus nerve promotes enterochromaffin cells to secrete serotonin (*70*), which in turn facilitates vagal function (*71–73*). Our results together with these previous findings prompted us to examine whether GFRAL^+^ neurons regulate intestinal type 2 immunity via vagal efferents.

We first examined whether there is a neural circuit connecting GFRAL^+^ neurons to the small intestine by polysynaptic retrograde tracing from the distal intestine (Methods). We injected the ileum with a pseudorabies virus (PRV) strain PRV-152 (fig. S8A), which resulted in robust labeling of neurons in the dorsal motor nucleus of the vagus nerve (DMV), NTS and AP (Fig. 5D; fig. S8B-D). Many GFRAL^+^ neurons in the NTS and AP were labeled by the PRV (Fig. 5D). Additional labeling was observed in several autonomic and limbic regions, including the PVN, lateral hypothalamus, medial CeA, PBN, and rostral ventrolateral medulla (RVLM) (fig. S8B-D). To further examine whether GFRAL^+^ neurons project to vagal preganglionic neurons in the DMV, we injected the AP of *Gfral^Cre^* mice with a Cre-dependent AAV expressing Synaptophysin-mRuby, followed by injecting the ileum of these mice with PRV-290 (fig. S9A; Methods). The mRuby-labeled presynaptic terminals originating from GFRAL^+^ neurons projected onto PRV-positive neurons in the DMV (fig. S9B). Finally, we conducted anterograde transsynaptic tracing of GFRAL^+^ neurons with a monosynaptic herpes simplex virus (HSV) (*74–76*) (fig. S9C; Methods). This assay revealed that GFRAL^+^ neurons make monosynaptic connections with DMV neurons expressing choline acetyltransferase (ChAT) (Fig. 5E; fig. S9D, E) — which are vagal preganglionic neurons — as well as with neurons in the NTS, nucleus ambiguus, PBN, PVN and CeA. Together, these results support the existence of a polysynaptic GFRAL^+^ neurons→DMV parasympathetic (vagal) neurons→intestine (GFRAL→DMV→intestine) pathway.

To determine whether the vagus nerve is required for the GFRAL^+^ neuron-driven intestinal responses, we performed subdiaphragmatic vagotomy (sdVx) or sham surgery (fig. S10A-C; Methods). Notably, sdVx prevented the increase in both tuft cells and goblet cells in the small intestine of NaChBac mice, such that the levels of tuft and goblet cells in NaChBac/sdVx mice were comparable to those in mice without NaChBac expression (both sham and sdVx) (Fig. 5F-I). SdVx did not prevent the body weight loss and the villus and crypt length reduction of NaChBac mice (fig. S10D-F), suggesting that these changes are mediated by circuitry other than the vagus nerve. Together, these results support a model in which GFRAL^+^ neurons regulate intestinal type 2 immunity through a polysynaptic vagal brain-gut circuit.

### GFRAL^+^ neurons protect against helminth infection

Helminth infection provides a physiologically relevant model to test the significance of GFRAL^+^ neuron function. In helminth infection, type 2 immunity orchestrates coordinated epithelial, immune and neuronal responses for host protection (*8, 9, 43, 44, 77–81*), promoting parasite clearance while limiting tissue damage and supporting tissue repair (*79, 80*). In particular, tuft cells release type 2 cytokines and ACh to stimulate mucus secretion by goblet cells, enhance chloride-dependent fluid flow and increase intestinal muscle contraction, collectively facilitating parasite expulsion in a “weep and sweep” response (*8, 82–84*).

Given our results that activation of GFRAL^+^ neurons is sufficient to drive hyperplasia of both tuft cells and goblet cells and increase type 2 cytokines, we reasoned that it might enhance host defense against helminths through the primed type 2 immune responses. To test this idea, we prepared a new cohort of NaChBac mice and their controls and infected them (s.c.) with the third-stage larvae (L3) of *Strongyloides venezuelensis* (*Sv*) (Fig. 6A; Methods), a well-characterized intestinal helminth that elicits type 2 immune responses and the canonical weep and sweep response (*79, 85, 86*). Parasite burden was quantified as fecal egg counts from day 5 to day 12 post-infection (Fig. 6B). Indeed, NaChBac mice exhibited reduced intestinal parasite burden on day 8 and 9, which was accompanied by an increase in plasma IgE antibodies (Fig. 6C). These results indicate that GFRAL^+^ neuron activity boosts intestinal type 2 immune defense to accelerate helminth clearance.

**Fig. 6.**
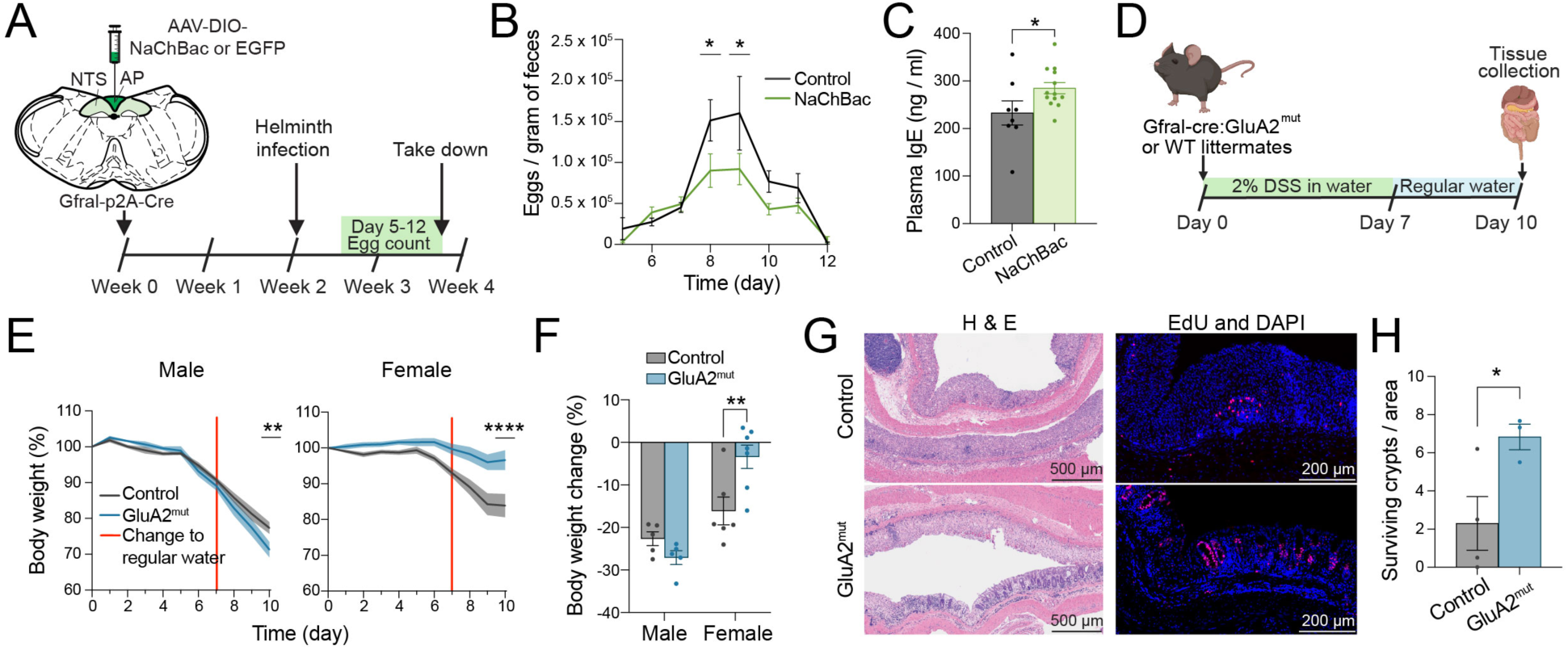
Enhanced type 2 immunity in the intestine following NaChBac activation leads to reduced S. venezuelensis burden. **(A)** Timeline of the experiments, in which NaChBac or eGFP was injected at week 0, followed by S. venezuelensis (Sv) infection at week 2. Fecal egg counts began 5 days after infection and continued until day 12, when mice were euthanized. **(B)** Quantification of fecal Sv eggs (Row factor:F (7, 133) = 18.79 and P<0.0001, Column factor: F (1, 19) = 5.886 and P=0.0254; day 8, *P = 0.0421; day 9, *P = 0.0160; two-way repeated-measures (RM) ANOVA followed by Sidak’s post hoc test). **(C)** Plasma IgE concentrations in NaChBac mice after Sv infection (*P = 0.0498; Unpaired t-test). **(D)** Timeline of the experiments, in which GluA2^mut^ or control littermates were treated with 2% DSS in drinking water for seven days, followed by a switch to regular drinking water on day 7 until day 10, when mice were euthanized. **(E)** Left: Male body weight over time (male; control, n = 5, GluA2^mut^, n = 5; Row factor: F (10, 80) = 181.7 and P<0.0001, Column factor: F (1, 8) = 0.8031 and P=0.3963; day 10, *P = 0.0065; two-way repeated-measures (RM) ANOVA followed by Sidak’s post hoc test). Right: Female body weight over time (female; control, n = 6, GluA2^mut^, n = 7; Row factor: F (10, 110) = 25.60 and P<0.0001, Column factor: F (1, 11) = 10.96 and P=0.0069; day 10, ****P < 0.0001; two-way repeated-measures (RM) ANOVA followed by Sidak’s post hoc test). **(F)** Body weight change from start to end in male and female mice (male, P = 0.2750, female, **P = 0.0017; Šídák’s multiple comparisons test**) (G)** (Left) Representative H&E images of colon tissue from GluA2^mut^ female mice; scale bar, 500 μm. (Right) Immunohistochemical images of the colon in GluA2^mut^ mice, showing EdU expression; scale bar, 200 μm. **(H)** Quantification of surviving crypts/area in female mice (control, n = 4, GluA2^mut^, n = 3; *P = 0.0495; Unpaired t test). Created in BioRender. van de Lisdonk, D. (2026) https://BioRender.com/dey02x5. Data are presented as mean s.e.m.

### GFRAL^+^ neurons protect female mice against DSS-induced colitis

Since type 2 immune responses promote tissue repair, they have been proposed as a treatment for colitis or IBD, during which tuft and goblet cell numbers are reduced, leading to increased epithelial permeability and disruption of mucus barrier (*6, 87*). Given that GFRAL^+^ neuron activation increased intestinal tuft cells (Fig. 3A-D; Fig. 5F-I), we reasoned that these neurons might protect against colitis (Fig. 6D; Methods).

To test this hypothesis, we adopted the dextran sulfate sodium (DSS)-induced colitis and used GluA2^mut^ mice for this model, because the NaChBac and DSS both resulted in marked weight loss (Fig. 6E; fig. S5H) and, if combined, they may lead to aggravated metabolic changes. We found that female GluA2^mut^ mice were protected against DSS-induced weight loss (Fig. 6E, F). Furthermore, epithelial proliferation was increased in the colon of these mice (Fig. 6G, H), consistent with the above results that GFRAL^+^ neuron activation promoted type 2-biased epithelial repair (Fig. 3A-D; Fig. 5F-I). In stark contrast, male GluA2^mut^ mice exhibited exacerbated DSS-induced weight loss (Fig. 6E, F), and several male GluA2^mut^ mice did not survive the DSS treatment (fig. S11A). It is possible that the male GluA2^mut^ mice’s lower baseline body weight made them vulnerable to the additional DSS-induced weight loss. Intestinal length did not differ between the treatment groups (fig. S11B), though male GluA2^mut^ mice displayed shorter colons than female GluA2^mut^ mice (fig. S11C). These results indicate a female-specific protective effect of the GluA2^mut^ against DSS-induced colitis. Together, these results demonstrate that activation of GFRAL^+^ neurons alters intestinal immunity and epithelial structure in a sex-dependent manner, which can enhance pathogen clearance or protect against inflammatory injury.

## Discussion

By integrating immune signals with the nutritional, metabolic, and other state signals, the brain coordinates local tissue defense with whole-body demands (*24, 26, 88–97*). Thus, the brain may function as “commander-in-chief” when whole-body needs call for enhanced protective immunity and epithelial repair, or restraint of excessive inflammation. The AP is particularly important in this process, as it is located outside of the BBB and thus is accessible by circulating factors (and with systemic therapeutic interventions) (*62, 98, 99*).

In this study, we investigated the role of AP GFRAL^+^ neurons in orchestrating type 2 immunity using mouse models of type 2-biased inflammation, combined with using mice in which the excitability of GFRAL^+^ neurons was chronically elevated. Fos mapping and electrophysiology experiments showed that both food allergen- and cytokine-driven type 2 inflammation activate GFRAL^+^ neurons. Furthermore, the inflammation condition leads to potentiated excitatory synaptic transmission onto GFRAL^+^ neurons. Circuit mapping and causal manipulation experiments demonstrated that GFRAL^+^ neurons regulate type 2 immunity via a vagal brain-gut circuit, shaping intestinal structure and pathophysiology in food allergy, helminth infection and colitis models. In parallel, these experiments also revealed that GFRAL^+^ neurons regulate water intake and control body weight independent of food intake and energy expenditure, and that many of the functions of GFRAL^+^ neurons are sex specific.

A striking consequence of chronically elevating GFRAL^+^ neuron excitability was the coordinated expansion of tuft and goblet cells and increased IL-4 and GDF15 expression in the intestine, hallmarks of a primed type 2 immune environment and characteristics of engagement of the canonical ‘weep and sweep’ program, in which tuft cell-derived signals and type 2 cytokines drive goblet cell hyperplasia, mucus secretion, luminal fluid flux and smooth muscle contraction to enhance barrier protection and parasite expulsion (*8, 9, 43, 77, 78, 100, 101*). The capacity of GFRAL^+^ neurons to elicit these features in the absence of active infection or inflammatory challenge underscores a role for central neural signals in preconditioning intestinal type 2 immunity. Indeed, promoting GFRAL^+^ neuron activity on one hand can accelerate parasite clearance during helminth infection, and on the other hand can facilitate epithelial regeneration in DSS-induced colitis. These results confirm the significance of GFRAL^+^ neurons in orchestrating the coordinated epithelial-immune response.

Mechanistically, GFRAL^+^ neuron function in type 2 immunity is mediated by a polysynaptic circuit composed of GFRAL^+^ neurons and postsynaptic DMV vagal preganglionic neurons projecting to the small intestine (GFRAL→DMV→intestine). GFRAL^+^ neuron activity causes elevated cholinergic and serotonergic signaling in the small intestine, likely through this circuit. The two signaling systems and their interactions might modulate type 2 immunity (*65–71*). Notably, we show that active type 2 immunity in turn led to enhanced activity of and potentiated excitatory synaptic transmission onto GFRAL^+^ neurons. Our results thus suggest the existence of an intestine→GFRAL→DMV→intestine feedback loop that amplifies type 2 immunity. Enterochromaffin cells (ECs) might play an important role in the feedback, as Tuft cell-derived acetylcholine drives EC serotonin release and vagal afferent activation during parasite-induced inflammation (*71*).

Acute activation of GFRAL^+^ neurons, with either chemogenetics or optogenetics, has been shown to suppress food intake, thereby causing weight loss (*29–32, 55–59*). We used GluA2^mut^- and NaChBac-based strategies to promote the excitability of these neurons, which also resulted in weight loss. However, this effect was mediated by a fundamentally different mechanism: despite that both the GluA2^mut^ mice and the NaChBac mice (particularly the latter) showed weight loss, their food intake remained unchanged. Instead, these mice had intestinal structural remodeling and reduced intestinal triglyceride content, pointing to impaired absorptive capacity as the primary driver of weight loss.

A possible reason why the GluA2^mut^ or NaChBac does not cause reduced food intake, as does chemogenetic or optogenetic activation of GFRAL^+^ neurons, is that the former strategies work more chronically than the latter, and that chronic activation of the GDF15/GFRAL system causes desensitization to anorexia. Supporting this view, it has been shown that, while GDF15 causes pregnancy-associated nausea and vomiting, pre-pregnancy levels of GDF15 predict the severity of these symptoms, with mothers genetically predisposed to high levels of GDF15 experiencing reduced pregnancy-associated nausea, whereas those predisposed to low levels of GDF15 having increased risk of developing hyperemesis gravidarum (*102, 103*). An open question is whether acute activation of GFRAL^+^ neurons primarily recruits aversive and anorexia-associated processes, while their slow and chronic elevation of activity leads to adaptive changes that uncouple these processes from metabolic and intestinal programs. Understanding this potential shift in circuit function could help develop strategies that promote beneficial metabolic effects while minimizing anorexia or aversive side effects.

In summary, we identify GFRAL^+^ neurons as a central neural regulator of intestinal type 2 immunity. By translating systemic immune signals into coordinated autonomic and possibly neuroendocrine outputs, these neurons promote type 2 immune remodeling, alters nutrient absorption and shapes disease outcomes. Our study reveals a previously unrecognized brain-gut-immune axis with potential therapeutic relevance for allergic, infectious and inflammatory diseases.

## Materials and Methods

### Mice

All animal experiments were approved by the Institutional Animal Care and Use Committee at Cold Spring Harbor Laboratory and adhered to the guidelines set by the U.S. National Institutes of Health. All experiments used male and female mice aged two months or older. Mice were group-housed (2-5 per cage) under a 12-hour light/dark cycle (lights on at 07:00, off at 19:00), with ad libitum access to food and water. All procedures were carried out during the light phase. Experimental and control groups consisted of littermates and were randomly assigned to each group. C57BL/6J mice (strain #000664) were obtained from Jackson Laboratory. Gfral-p2A-Cre mice, originally developed by Stephen Liberles (*57*), were bred with heterozygous Rosa26-stopflox-GluA2^mut^ mice, generated as previously described (*46*), to produce Gfral-p2a-Cre:GluA2^mut^ offspring. GluA2^mut^ mice contain two-point mutations in the AMPA receptor GluA2 subunit expressed in a Cre-dependent manner. The L483Y mutation impairs AMPA receptor desensitization (*47, 48*), whereas the R845A mutation prevents AMPA receptor endocytosis (*49*). Together, these mutations enhance excitatory synaptic transmission. Gfral-p2A-Cre positive littermates lacking the GluA2^mut^ allele were used as controls. All mouse lines were maintained on a C57BL/6J genetic background.

### Disease models and immune stimulation paradigms Food allergy challenge

Mice were subcutaneously sensitized on days 0 and 7 with endotoxin-free OVA (0.25 mg/kg; BioVendor, 321001) mixed with alum adjuvant (50 mg/kg; Invivogen, vac-alu-250) and brought to a final injection volume of 0.2 mL in PBS. Control animals received the alum + PBS preparation without OVA. To acclimate mice to oral gavage before allergen exposure, all animals were administered 0.2 mL of regular drinking water by gavage on days 10 and 12. Allergen challenges were performed on days 14 and 16 by delivering 50 mg of grade III OVA (Sigma-Aldrich, A5378) dissolved in 0.2 mL of drinking water.

### IL25 challenge

Mice were administered daily IP injections for seven consecutive days, receiving either 0.2 mL of 500 ng recombinant IL25 (R&D systems, 1399-IL-025/CF) or 0.2 mL of PBS as a control. Mice were euthanized 6 hours after the final injection to assess Fos expression in the brain.

### Helminths

For the helminth infection model, mice were infected subcutaneously with 700 Strongyloides venezuelensis (Sv) L3 larvae in 200 µL of water. Mice were euthanized on day 12 post-infection. Beginning on day 5 post-infection, fecal pellets were collected daily through day 12 to quantify parasite eggs burdens.

### Induction of DSS colitis

For the induction of colonic inflammation (colitis), GluA2^mut^ mice and their WT littermates were administered 2% dextran sulfate sodium (DSS; MP Biomedicals, 9011-18-1) in their drinking water for 7 days, followed by 3 days of recovery with regular drinking water. Body weight was recorded daily at 10:00. Four hours before euthanasia, mice were injected intraperitoneally with 5-ethynyl-2’-deoxyuridine (EdU, 0.1 mg; Invitrogen, C10640), following the manufacturer’s recommendations, to assess colonic epithelial cell proliferation and migration by specific staining. Surviving crypts per area were determined based on EdU-positive crypts in an area of the colon that was significantly damaged; the analyzed ‘area’ was 850 µm width.

### Surgical and neural manipulation procedures

#### Viruses

The pAAV-EF1a-DIO-2a-mNaChBac was produced by Baylor College of Medicine. The pAAV-hSyn-FLEX-TeLC-P2A-EYFP-WPRE, pAAV-hSyn-DIO-EGFP (AAV5) and pAAV hSyn FLEx mGFP-2A-Synaptophysin-mRuby (AAV1) were purchased from Addgene (Watertown, MA 02472, USA). PRV-152 and PRV-290 was purchased from Center for Neuroanatomy with Neurotropic Viruses (CNNV; P40 OD010996)). The AAV9-CAG-FLEX-TeLC-GFP was made by the Penn Vector Core (Philadelphia, PA). The rAAV-EFla-DIO-EGFP-T2A-TK-WPRE-hGH polyA and HSVΔTK-hUbC-tdTomato were purchased from BrainVTA. Custom viral particles were produced using this vector by WZ Biosciences Inc. (Rockville, MD 20855, USA). All viruses were aliquoted and stored at -80 °C until use.

#### Stereotaxic surgery

Stereotaxic injections were carried out following standard surgical protocols, as detailed in prior studies (*104, 105*). All procedures were conducted under sterile conditions, and body temperature was maintained at 35°C using a heating pad throughout the surgery. Anesthesia was induced and maintained with 1-2% isoflurane, and animals received preemptive analgesia and anti-inflammatory treatment with Metacam (5 mg/kg) prior to the procedure. Mice were placed in a stereotaxic injection frame (Angle Two Stereotaxic System, myNeuroLab.com) equipped with a digital brain atlas interface to assist in accurately locating and targeting brain regions. The following coordinates were used for AP: 7.65 mm posterior to bregma, 0.0 mm lateral to midline, and 4.66 mm below the skull surface. A small incision was made to expose the skull, and a 1 mm diameter craniotomy was performed at the injection site. Virus (0.1-0.3 μL) was delivered via a glass micropipette (tip diameter: ∼5 μm) using a pressure injection system (5-20 psi, 5-20 ms pulses at 0.5 Hz), controlled by a Picospritzer III (General ValveParker, Hollis, NH, USA) and a pulse generator (Agilent). Following the injection, animals received subcutaneous Metacam (1-2 mg/kg meloxicam) for postoperative pain management and were returned to their home cages for monitoring during recovery.

#### Subdiaphragmatic vagotomy (sdVx)

Mice were anesthetized with 1-2% isoflurane and prepared for surgery by removing abdominal hair and disinfecting the area. To minimize pain and inflammation, animals were pretreated with Metacam (5 mg/kg) before the procedure. They were then placed on a sterile operating surface over a heating pad set to 35°C and draped with a sterile cover to maintain aseptic conditions. A longitudinal incision was made through the abdominal skin and peritoneum to expose the abdominal cavity. The liver was carefully lifted and stabilized using saline-moistened cotton swabs to allow visualization of the lower esophagus. Using a surgical microscope, the right and left vagus nerves running along the esophagus just below the diaphragm were identified. Both nerves were cut proximally using fine micro-scissors. In sham-operated mice, the vagus nerves were exposed but left intact. To assess whether GFRAL activation signals to the intestine via the vagus nerve, a subset of mice underwent stereotaxic surgery following vagotomy. pAAV-EF1a-DIO-2A-mNaChBac was stereotaxically injected into the AP using the surgical procedures described above. Postoperatively, animals received subcutaneous Metacam (1-2 mg/kg) for analgesia and were returned to their home cages for recovery and monitoring. To confirm the vagatomy, 200 µL of 4 mg/mL Fluoro-Gold was administered IP four days prior to tissue collection, successful vagotomy was confirmed by the absence of dye labeling in the dorsal motor nucleus of the vagus.

#### PRV tracing

Mice were anesthetized with 1-2% isoflurane and prepared for surgery by shaving and disinfecting the abdominal region. Preemptive analgesia and anti-inflammatory treatment were administered with Metacam (5 mg/kg) prior to the procedure. Animals were positioned on a sterile surgical platform over a heating pad maintained at 35°C, and the operative area was covered with a sterile drape to maintain aseptic conditions. A midline incision was made through the skin and underlying peritoneum to expose the abdominal cavity. For small intestine injections, the distal portion of the intestine was carefully exteriorized and stabilized using two sterile swabs. A total of 2 µL of PRV-152 or PRV-290 was injected using a pulled glass micropipette attached to a Nanoliter 2010 injector (WPI). After injections, the muscle layer was closed with absorbable sutures, and the skin was closed using staples. Postoperative analgesia was provided with subcutaneous Metacam (1-2 mg/kg), and animals were returned to their home cages for recovery and monitoring. Intestinal and brain tissues were collected 4-5 days following PRV-152 injection for downstream analyses.

#### In vitro electrophysiology

Acute brain slices were prepared from mice aged 2 to 4 months. Mice were anaesthetized with 4% isoflurane and euthanized by rapid decapitation. Brain was rapidly removed, and chilled in ice-cold dissection buffer (composed of (in mM): 110.0 Choline chloride, 25.0 NaHCO3, 1.25 NaH2PO4, 2.5 KCl, 25.0 glucose, 0.5 CaCl2, 7.0 MgCl2, 11.6 ascorbic acid, and 3.1 pyruvic acid, bubbled with 95% O_2_ and 5% _CO2_). Coronal sections (250-300 μm thick) containing the AP were cut using a HM650 Vibrating-blade Microtome (Thermo Fisher Scientific). Slices were transferred to a recovery chamber containing artificial cerebrospinal fluid (ACSF) solution (containing (in mM): 118 NaCl, 2.5 KCl, 26.2 NaHCO3, 1 NaH2PO4, 20 Glucose, 2 CaCl2 and 2 MgCl2, pH 7.4, bubbled with 95% O_2_ and 5% CO_2_). The slices were incubated at 34°C for at least 40 minutes and then held at room temperature (20-24°C) until use. Electrophysiological recordings were made in ACSF under continuous flow, bubbled with 95% O₂ and 5% CO₂. Whole-cell patch-clamp recordings from AP neurons were obtained using a Multiclamp 700B amplifier and pCLAMP 10 software (Molecular Devices, Sunnyvale, CA). Recordings were guided with an Olympus BX51 microscope equipped with both transmitted light and epifluorescence illumination (Olympus Corporation, Tokyo, Japan). GFRAL^+^ neurons were identified based on the presence of red fluorescence in Gfral-p2A-Cre:Ai14 mice or green fluorescence in Gfral-p2A-Cre:GluA2^mut^ 2(Y845A) mice. Synaptic activity was recorded at a holding potential of -70 mV to isolate AMPA receptor-mediated currents, with signals low-pass filtered at 1 kHz. The internal solution used for voltage-clamp recordings contained (in mM) 115 Cesium methanesulfonate, 20 CsCl, 10 HEPES, 2.5 MgCl2, 4 Na2-ATP, 0.4 Na3-GTP, 10 Na-phosphocreatine, and 0.6 EGTA, to pH 7.2. Miniature excitatory post-synaptic currents (mEPSCs) were recorded in the presence of 1 μM tetrodotoxin (TTX) and 100 μM picrotoxin (PTX). mEPSC data were analyzed using an automated machine learning tool (*106, 107*).

### Tissue processing and histology

#### Immunohistochemistry

Mice were anesthetized using isoflurane and Euthasol (0.1 mL; Virbac, Fort Worth, TX, USA), then transcardially perfused with 30 mL of 1x phosphate-buffered saline (PBS), followed by 30 mL of 4% paraformaldehyde (PFA) in 1x PBS. Brains were carefully removed and placed in 4% PFA for overnight post-fixation at room temperature (RT) on a shaker. The following day, tissues were transferred to a 30% sucrose solution in PBS for cryoprotection, incubated overnight at RT on a shaker. Coronal brain slices (50 μm thick) were prepared using a freezing microtome (Leica SM 2010R, Leica). Sections were incubated in a blocking solution containing 5% bovine serum albumin (BSA) and 0.1% Triton X-100 in PBS for 1 hour at RT with shaking. Primary antibody was added and the sections were incubated overnight at 4°C. Sections were washed (3 x 15 minutes) in PBS and then incubated with secondary antibodies for 2 hours at RT. Following another round of PBS washes (3 x 15 minutes), the sections were mounted onto glass slides using Fluoromount-G (eBioscience, San Diego, CA, USA). Images were taken with a LSM 710 or LSM 980 laser-scanning confocal microscope (Carl Zeiss, Oberkochen, Germany). Primary antibodies used were rabbit anti-cFOS (1:500, Cell Signaling, 2250S; 1:500 Synaptic systems, #226308), sheep anti-Gfral (1:200, R&D systems, #AF5728), chicken anti-GFP (1:1000, AvesLabs, #GFP-1020), rabbit anti-PKCd (1:500, Abcam, #ab182126), guinea pig anti-cFOS (1:1000, Synaptic systems, #226308), goat anti-ChAT (1:200, Sigma-Aldrich, #AB144P), rabbit anti-RFP (1:1000, Rockland, #600-401-379). The fluorophore-conjugated secondary antibodies used were Alexa Fluor-594 donkey anti-rabbit IgG (1:500-1:1000, Life technologies #a21207), Alexa Fluor-Cy3 donkey anti-sheep IgG (1:500, Jackson ImmunoResearch, #713-165-003), Alexa Fluor-488 donkey anti-chicken IgG (1:1000, Jackson ImmunoResearch, #703-545-155), Alexa Fluor-647 donkey anti-guinea pig IgG (1:1000, Jackon ImmunoResearch, #706-605-148), Alexa Fluor-647 donkey anti-goat IgG (1:500, Invitrogen, #A32849). DAPI (0.5 μg/ml in PBS, Invitrogen, D1306) was used to stain nuclei.

#### FFPE processing and staining

Muscle, inguinal white adipose tissue (iWAT), and brown adipose tissue (BAT) were collected and weighed. The small intestine and colon were harvested, carefully stripped of attached fat, and opened lengthwise before being rinsed with ice-cold PBS to remove any luminal contents. The small intestine was separated into three segments, duodenum, proximal jejunum, and ileum, and each portion was rolled in a “Swiss-roll” configuration, beginning with the distal end at the core and the proximal end on the outside, maintaining the natural spatial orientation of the tissue. All tissues were fixed in 4% PFA for subsequent histological analyses. Formalin-fixed tissues were processed and infiltrated with paraffin using a Thermo Excelsior ES processor, then embedded with a Thermo HistoStar system following standard protocols. Embedded blocks were cut into 10 µm sections and mounted onto slides (VWR Superfrost Plus, 48311-703). H&E staining was carried out at the CSHL tissue imaging facility using a Leica ST5020 Multistainer. Sections were first deparaffinized and rehydrated, then immersed in hematoxylin (Hematoxylin 560 MX, Leica) for 1 minute, rinsed in Define MX-aq (Leica) for 30 seconds, and treated with Blue Buffer 8 (Leica) for 1 minute to achieve bluing. Slides were then stained in eosin (Eosin 515 LT, Leica) for 30 seconds, dehydrated through graded alcohols, and finally cover slipped using a Leica CV5030 automated cover slipper.

For immunofluorescence, FFPE slides were deparaffinized in xylene (3 x 5 min) and rehydrated through graded alcohols (3 x 100%, 95%, and 70%, 5 min each). Antigen retrieval was performed for 6 min in 1 mM citrate buffer (Sigma-Aldrich, C9999). Sections were then blocked with PBS containing 1% BSA and 10% normal donkey serum and incubated with primary antibodies. Primary antibodies included rabbit anti-Dclk1 (1:300, Cell Signaling, #62257S), goat anti-E-cadherin (1:300, R&D Systems, #AF748), and rabbit anti-Muc2 (1:500, Abcam, #ab272692). Fluorophore-conjugated secondary antibodies were Alexa Fluor 594 donkey anti-rabbit IgG (1:500, Life Technologies, #A21207) and Alexa Fluor 647 donkey anti-goat IgG (1:500, Invitrogen, #A32849). Nuclei were counterstained with DAPI (0.5 µg/mL in PBS, Invitrogen, D1306). Stained sections were imaged using an Olympus VS200 Slide Scanner.

#### LifeCanvas tissue clearing

Whole-brain tissue clearing and immunofluorescence were carried out using the SHIELD preservation method as previously described (*108*). In brief, mice were transcardially perfused with 30 mL of 1x PBS followed by 4% PFA in PBS. Brains were removed, immersed in 4% PFA at 4°C for 24 hours on a shaker, and subsequently rinsed in 1x PBS. Post-fixation preservation was performed using SHIELD reagents (LifeCanvas Technologies, SH-250), and tissues were then subjected to delipidation with Clear+ delipidation reagents (LifeCanvas Technologies, DB) according to established protocols (*108*). Following delipidation, brains were incubated in 50% EasyIndex (refractive index = 1.52; LifeCanvas Technologies, EI-100) for 24 hours at 37°C and then transferred to 100% EasyIndex for an additional 24 hours to match refractive index. After index matching, samples were embedded in 2% agarose (Sigma-Aldrich, A5030) prepared in EasyIndex. Cleared and embedded brain were imaged using a SmartSPIM axially swept light-sheet microscope equipped with a 3.6 x objective (0.2 NA). Three-dimensional image stacks generated by the light-sheet were processed and analyzed using ImageJ.

### Molecular and biochemical assays

#### RNA isolations and qPCR

Sections of small intestine were collected into ZR BashingBead Lysis Tubes (Zymo Research, S6003-50) containing TRIzol and homogenized using a bead-based tissue disruptor (Benchmark Scientific, D1036). RNA extraction was performed with the Direct-zol RNA Microprep Kit (Zymo Research, R2061) according to the manufacturer’s instructions. For cDNA synthesis, 500 ng of total RNA was reverse-transcribed using the SuperScript™ IV VILO™ Master Mix (Invitrogen, 11756050). Quantitative RT-PCR was carried out on a QuantStudio 3 instrument (Applied Biosystems) using TaqMan Fast Advanced Master Mix (Applied Biosystems, 4444557). TaqMan probes targeting IL4 (Life technologies, 4453320/Mm00445259_m1), IL5 (Life technologies, 4453320/Mm00439646_m1), IL13 (Life technologies, 4453320/Mm00434204_m1), GDF15 (Life technologies, 4331182/Mm00442228_m1) and Chrna7 (4453320/Mm01312230_m1) were used. GAPDH (Life technologies 4331182/Mm99999915_g1) was used as the endogenous control. Relative gene expression was quantified using the 2-ΔΔCt method.

#### Triglyceride measurements

Intestinal triglycerides were quantified using a commercially available triglyceride assay kit (Abcam, ab65336). Approximately 30 mg of small intestinal tissue was combined with 0.5 mL of 5% NP-40 and mechanically disrupted. The resulting lysate was heated to 95 °C for 5 minutes, cooled, and subjected to a second identical heating step to ensure complete solubilization. Samples were then brought to room temperature and centrifuged at 15,000 RPM for 2 minutes, after which the clarified supernatant was collected for measurement. For the assay, 50 µL of each supernatant was loaded into the designated wells of the assay plate and processed according to the manufacturer’s protocol. Triglyceride content was determined by converting measured concentrations to total mass based on initial tissue weight and subsequently normalizing values to total protein content. Protein levels in the lysates were determined using the bicinchoninic acid (BCA) assay (Thermo Scientific, A55865). Readouts for triglyceride and protein assays were obtained using microplate readers (SpectraMax i3 MiniMax 300; Molecular devices), and data acquisition was performed with Softmax Pro (v6.3) software.

#### Plasma measurements

Blood samples were collected into heparin-coated tubes to prevent clotting. Tubes were spun at 2000 g for 20 minutes at 4°C, after which plasma was separated, aliquoted and rapidly frozen in liquid nitrogen before storage at -80°C. Plasma concentrations of GDF-15 and IgE were quantified using commercially available ELISA kits: Mouse/Rat GDF-15 Quantikine ELISA Kit (R&D systems; MGD150) and the Mouse IgE ELISA kit (ABCAM; ab157718).

#### Fecal Water Content

Pre-weighed tubes were used to collect 2-3 fecal pellets from each mouse, and the wet weight was measured. Tubes were then placed in an oven at 60 °C overnight to dry the samples. The dry weight of the fecal pellets was measured, and the water content was calculated as the difference between wet and dry weight, normalized to the wet weight: (wet weight – dry weight) / wet weight.

#### Small intestine neurotransmitters targeted LC-MS/MS

Sample extraction process: The sample was thawed on ice. 50 mg (± 2.5 mg) of the sample was added to 500 μL of pre-chilled 70% methanol/water extraction solution and vortexed for 3 minutes. The mixture was centrifuged at 12,000 r/min for 10 minutes at 4°C. 300 μL of the supernatant was transferred to a new tube, incubated at -20°C for 30 minutes, and then centrifuged at 12,000 r/min for 10 minutes at 4°C. 200 μL of the supernatant was collected for LC-MS/MS analysis.

Chromatography - Mass Spectrometry Acquisition Conditions: UPLC Conditions: The sample extracts were analyzed using an LC-ESI-MS/MS system (UPLC, ExionLC AD, https://sciex.com.cn /; MS, QTRAP® 6500+ System, https://sciex.com /). The analytical conditions were as follows, HPLC: column, Waters ACQUITY UPLC HSS T3 C18 (100 mm×2.1 mm i.d.,1.8 µm); solvent system, water with 0.1% formic acid (A), acetonitrile with 0.1% formic acid (B); The gradient was started at 5% B (0 min), increased to 95% B (0-8 min), 95% B (8-9.5 min), finaly ramped back to 5% B (9.6-12 min); flow rate, 0.35 mL/min; temperature, 40°C; injection volume: 2 μL.

ESI-MS/MS Conditions: AB 6500+ QTRAP® LC-MS/MS System, equipped with an ESI Turbo Ion-Spray interface, operating in both positive and negative ion modes and controlled by Analyst 1.6 software (AB Sciex). The ESI source operation parameters were as follows: ion source, turbo spray; source temperature 550°C; ion spray voltage (IS) 5500 V(Positive), -4500 V(Negative); curtain gas (CUR) were set at 35.0 psi; DP and CE for individual MRM transitions was done with further DP and CE optimization. A specific set of MRM transitions were monitored for each period according to the neurotransmitters eluted within this period.

Qualitative and Quantitative Principles of Metabolites: Metabolites were quantified by multiple reaction monitoring (MRM) using triple quadrupole mass spectrometry. In MRM mode, the first quadrupole screened the precursor ions for the target metabolite and excluded ions of other molecular weights. After ionization induced by the impact chamber, the precursor ions were fragmented, and a characteristic fragment ion was selected through the third quadrupole to exclude the interference of non-target ions. After obtaining the metabolite spectrum data from different samples, the peak area was calculated on the mass spectrum peaks of all metabolite and analyzed by standard curves.

### Metabolic cages

Mice were housed individually and given one week to acclimate to the metabolic monitoring system (CLAMS; Columbus Instruments) before data collection. All recordings were obtained under a standard 12-hour light/dark cycle (lights on from 06:00 to 18:00). The CLAMS system continuously monitored food and water consumption, respiratory exchange ratio (RER), energy expenditure, oxygen consumption (VO₂), carbon dioxide production (VCO₂), and locomotor activity. Animals were maintained on a standard chow diet (PicoLab Rodent Diet 20, #5053*; 3.43 kcal/g) throughout the experiment. Data acquisition was performed using CLAX software (Columbus Instruments), and downstream analyses were carried out using customized MATLAB scripts (*109*). Metabolic variables were examined across a 72-hour interval and presented either as mean values or as time-series over the 72 hours.

### Data analysis and statistics

Statistical analyses were carried out using GraphPad Prism 10 (GraphPad Software, La Jolla, CA), MATLAB (MathWorks, Natick, MA) and RStudio (Posit PBC, Boston, MA). Data are presented as mean and error bars represent ± SEM. Normality of each dataset was assessed with the Shapiro-Wilk normality test. Depending on the distribution, either unpaired t-tests (for normally distributed data) or Mann-Whitney U test (for non-normal data) were applied. To detect differences between three unpaired groups, the one-way ANOVA followed by Sidak’s multiple comparisons test was used. For comparisons involving two independent groups, (repeated measures (RM)) two-way ANOVA followed by Sidak’s multiple comparisons post hoc testing was used. Statistical significance was defined as p < 0.05 for all analyses.

## Acknowledgments

We thank Daniel Mucid for providing the Strongyloides venezuelensis, members of the Li laboratory for helpful discussions, Jill Habel for assistance with experiments using the metabolic cages, Nikolas Holland for assistance with the subdiaphragmatic vagotomy surgery, and Innomics for assistance with the small intestine neurotransmitters targeted LC-MS/MS. We thank the Animal Core, Microscopy Core, Histology Core, Neuroimaging and Behavior Core of Cold Spring Harbor Laboratory (CSHL) for their help.

## Funding

CSHL and Northwell Health Affiliation

The Robert Lindsay Foundation

Key R&D Program of Zhejiang 2024SSYS0031

## Author contributions

Conceptualization: D.V.D.L., O.E., B.L.

Methodology: D.V.D.L., O.E., B.L.

Investigation: D.V.D.L., O.E., Z.K., W.G.

Visualization: D.V.D.L.,

Funding acquisition: B.L.

Project administration: B.L.

Supervision: S.B., D.M., H.K., B.L.

Writing – original draft: D.V.D.L., B.L.

Writing – review & editing: D.V.D.L., B.L.

## Data, code, and materials availability

All data supporting this study are available from the corresponding author upon request. All other data are available in the manuscript or the supplementary materials.

## FIGURES and SUPPLEMENTARY FIGURES

**Fig. S1.**
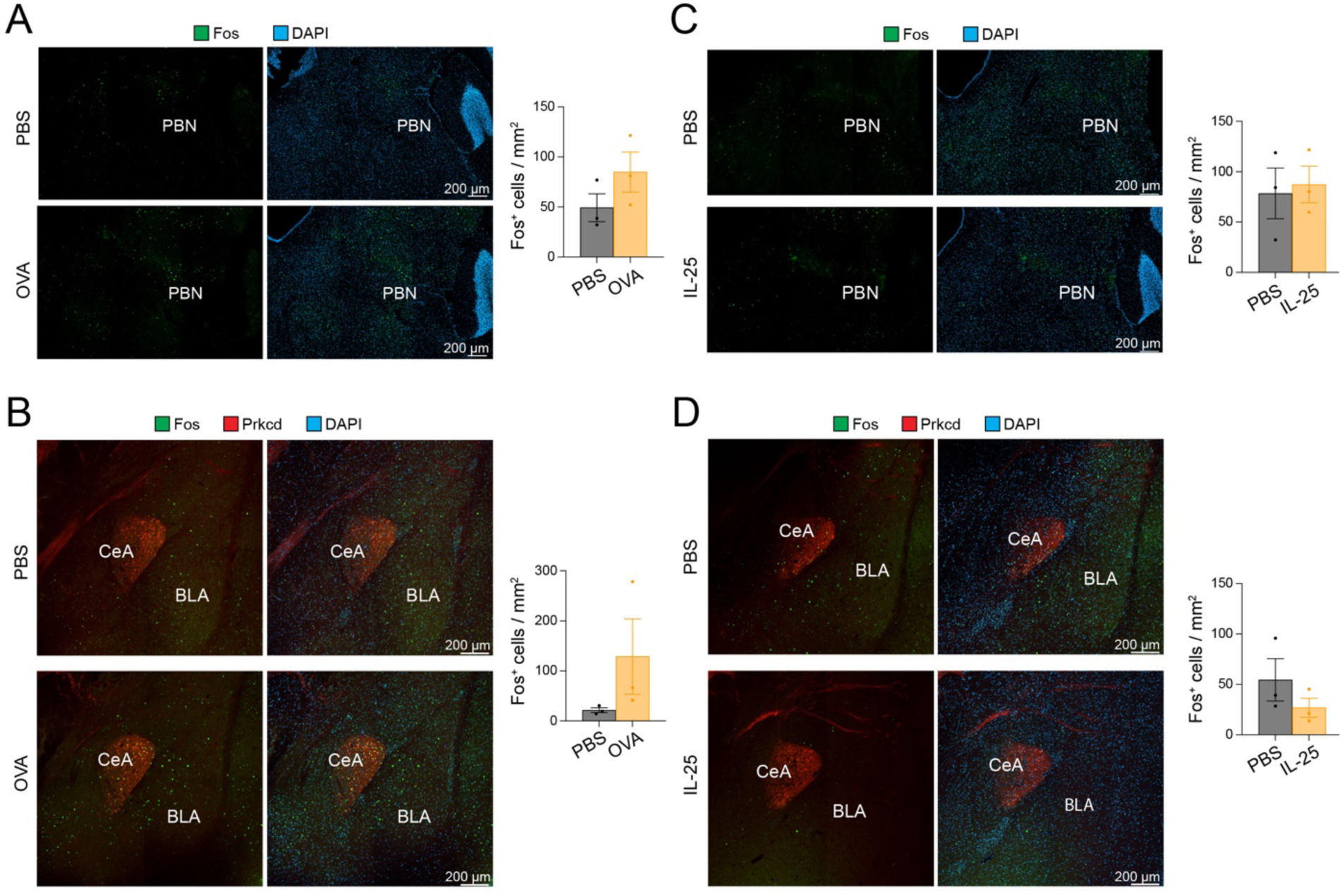
Fos expression across different brain areas following type 2 immune responses. **(A)** Immunohistochemical images of the PBN stained for Fos after OVA gavage; scale bar, 200 μm (PBS + alum n = 3, OVA + alum n = 3) (P = 0.2183; Unpaired t test). **(B)** Immunohistochemical images of the CeA stained for Fos after OVA gavage; scale bar, 200 μm (P = 0.2272; Unpaired t test). **(C)** Immunohistochemical images of the PBN stained for Fos after IL-25 or PBS treatment; scale bar, 200 μm (PBS n = 3, IL25 n = 3) (P = 0.7904; Unpaired t test). **(D)** Immunohistochemical images of the CeA stained for Fos after IL-25 or PBS treatment; scale bar, 200 μm (P = 0.2939; Unpaired t test). PBN, parabrachial nucleus and CeA, central amygdala, BLA, basolateral amygdala. Data are presented as mean s.e.m.

**Fig. S2.**
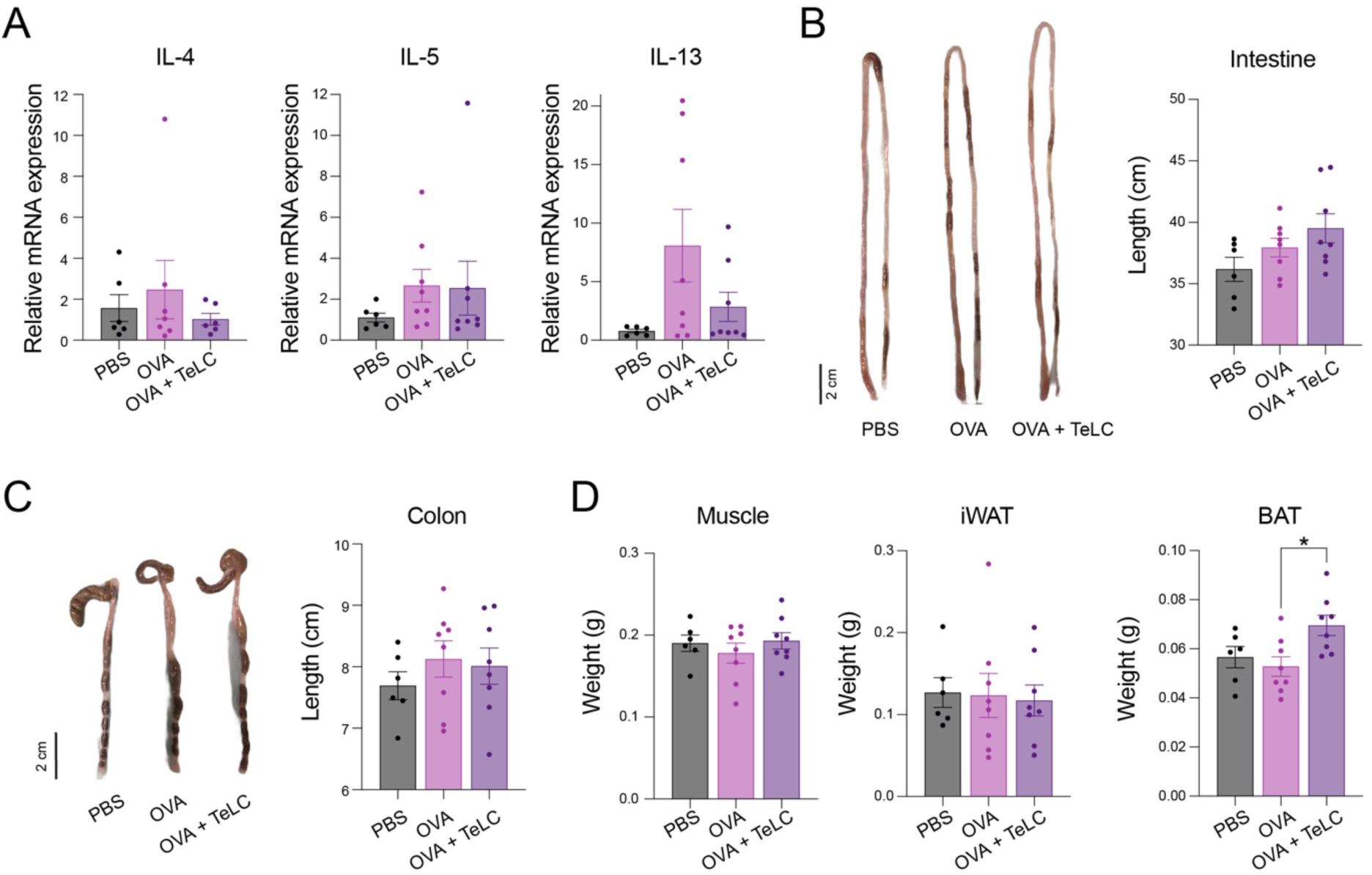
Inhibition of GFRAL neurons does not significantly affect type 2 cytokine responses during food allergy. **(A)** Expression of IL-4, IL-5 and IL-13 mRNA in the small intestine of TeLC mice (IL-4, PBS + alum (n = 6) vs. OVA + alum (n = 7), P = 0.7971; PBS + alum vs. OVA + alum + TeLC (n = 6), P = 0.9246; OVA + alum vs. OVA + alum + TeLC, P = 0.5641) (IL-5, PBS + alum (n = 6) vs. OVA + alum (n = 8), P = 0.5319; PBS + alum vs. OVA + alum + TeLC (n = 8), P = 0.5834; OVA + alum vs. OVA + alum + TeLC, P = 0.9953) (IL-13, PBS + alum (n = 6) vs. OVA + alum (n = 8), P = 0.0873; PBS + alum vs. OVA + alum + TeLC (n = 8), P = 0.8860; OVA + alum vs. OVA + alum + TeLC, P = 0.2340; one-way ANOVA followed by Šídák’s multiple comparisons test). **(B)** Representative intestine (left) and intestinal length (right) in control, GFP with OVA + alum, and TeLC with OVA + alum treated mice (PBS+alum vs OVA+alum, P = 0.5678; PBS+alum vs OVA+alum+TeLC, P = 0.1002; OVA+alum vs OVA+alum+TeLC, P = 0.5957; one-way ANOVA followed by Sidak’s post hoc test). **(C)** Representative colon (left) and colon length (right) in control, GFP with OVA + alum, and TeLC with OVA + alum treated mice (PBS+alum vs OVA+alum, P = 0.6724; PBS+alum vs OVA+alum+TeLC, P = 0.8380; OVA+alum vs OVA+alum+TeLC, P = 0.9873; one-way ANOVA followed by Sidak’s post hoc test). **(D)** Weights of skeletal muscle and adipose tissue (muscle; PBS + alum vs. OVA + alum, P = 0.8511; PBS + alum vs. OVA + alum + TeLC, P = 0.9970; OVA + alum vs. OVA + alum + TeLC, P = 0.7026) (iWAT; PBS + alum vs. OVA + alum, P = 0.9993; PBS + alum vs. OVA + alum + TeLC, P = 0.9879; OVA + alum vs. OVA + alum + TeLC, P = 0.9962) (BAT; PBS + alum vs. OVA + alum, P = 0.9001; PBS + alum vs. OVA + alum + TeLC, P = 0.1310; OVA + alum vs. OVA + alum + TeLC, *P = 0.0224; one-way ANOVA followed by Sidak’s post hoc test). Data are presented as mean s.e.m.

**Fig. S3.**
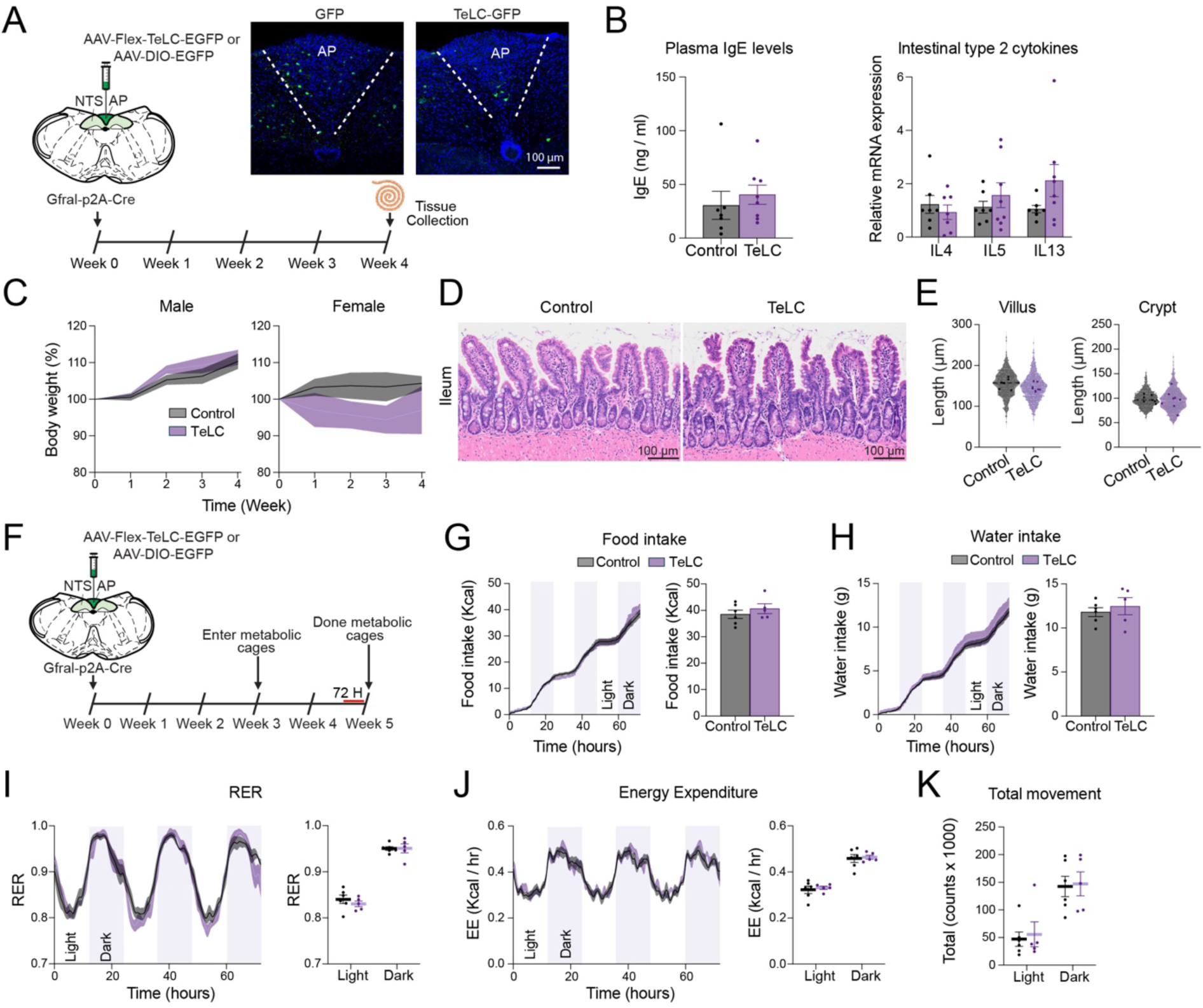
Metabolic effects of GFRAL neuron inhibition. **(A)** Schematic of TeLC or eGFP injection in AP. Example immunohistochemical images of the AP showing expression of eGFP or TeLC; scale bar, 100 μm. **(B)** Left, plasma IgE concentrations in control and TeLC mice (control, n = 7, TeLC, n = 8; P = 0.3357; Mann-Whitney test). Right, expression of IL-4, IL-5 and IL-13 mRNA in the small intestine of (left) male and (right) female TeLC mice (IL-4 control, n = 7, TeLC, n = 7; P = 0.9384; IL-5 control, n = 7, TeLC, n = 8; P = 0.8114; IL-13 control, n = 7, TeLC n = 8; P = 0.1698; Šídák’s multiple comparisons test). **(C)** Left, male body weight over time (male; control, n = 4, TeLC, n = 4; Row factor: F (1.276, 7.658) = 48.87 and P<0.0001, Column factor: F (1, 6) = 1.246 and P=0.3070; week 1, P = 0.9656; week 2, P = 0.5754; week 3, P = 0.8907; week 4, P = 0.9355; two-way repeated-measures (RM) ANOVA followed by Sidak’s post hoc test). Right, female body weight over time (female; control, n = 3, TeLC, n = 4; Row factor: F (1.937, 9.684) = 0.1357 and P=0.8688, Column factor: F (1, 5) = 1.733 and P=0.2451; week 1, P = 0.8255; week 2, P =0.6373; week 3, P = 0.6035; week 4, P = 0.8819; two-way repeated-measures (RM) ANOVA followed by Sidak’s post hoc test). **(D)** Representative HE images from Ileum in TeLC mice; scale bar, 100 μm. **(E)** Quantification of villus and crypt length (control, n = 7, TeLC, n = 8; Villus P = 0.0917; Crypt, P = 0.8176; Unpaired t test). **(F)** Timeline of the metabolic cage experiments in TeLC mice with the last 72 hours analyzed from the final week in the metabolic cages. **(G)** Food intake in TeLC mice over time (Left) and average food intake (Right) (P = 0.4027; Unpaired t test). **(H)** Water intake in TeLC mice over time (Left) and average water intake (Right) (P = 0.5343; Unpaired t test). **(I)** Respiratory exchange ratio (RER) in TeLC mice over time (Left) and average RER (Right) (control, n = 6, TeLC, n = 5; Light, P = 0.5992, Dark, P = 0.9996; Šídák’s multiple comparisons test). **(J)** Energy expenditure (EE) in TeLC mice over time (Left) and average EE (Right) (Light, P = 0.8987, Dark, P = 0.9569; Šídák’s multiple comparisons test). **(K)** Total movement in TeLC mice (Light, P = 0.9379, Dark, P = 0.9793; Šídák’s multiple comparisons test). Created in BioRender. van de Lisdonk, D. (2026) https://BioRender.com/dey02x5. Data are presented as mean s.e.m.

**Fig. S4.**
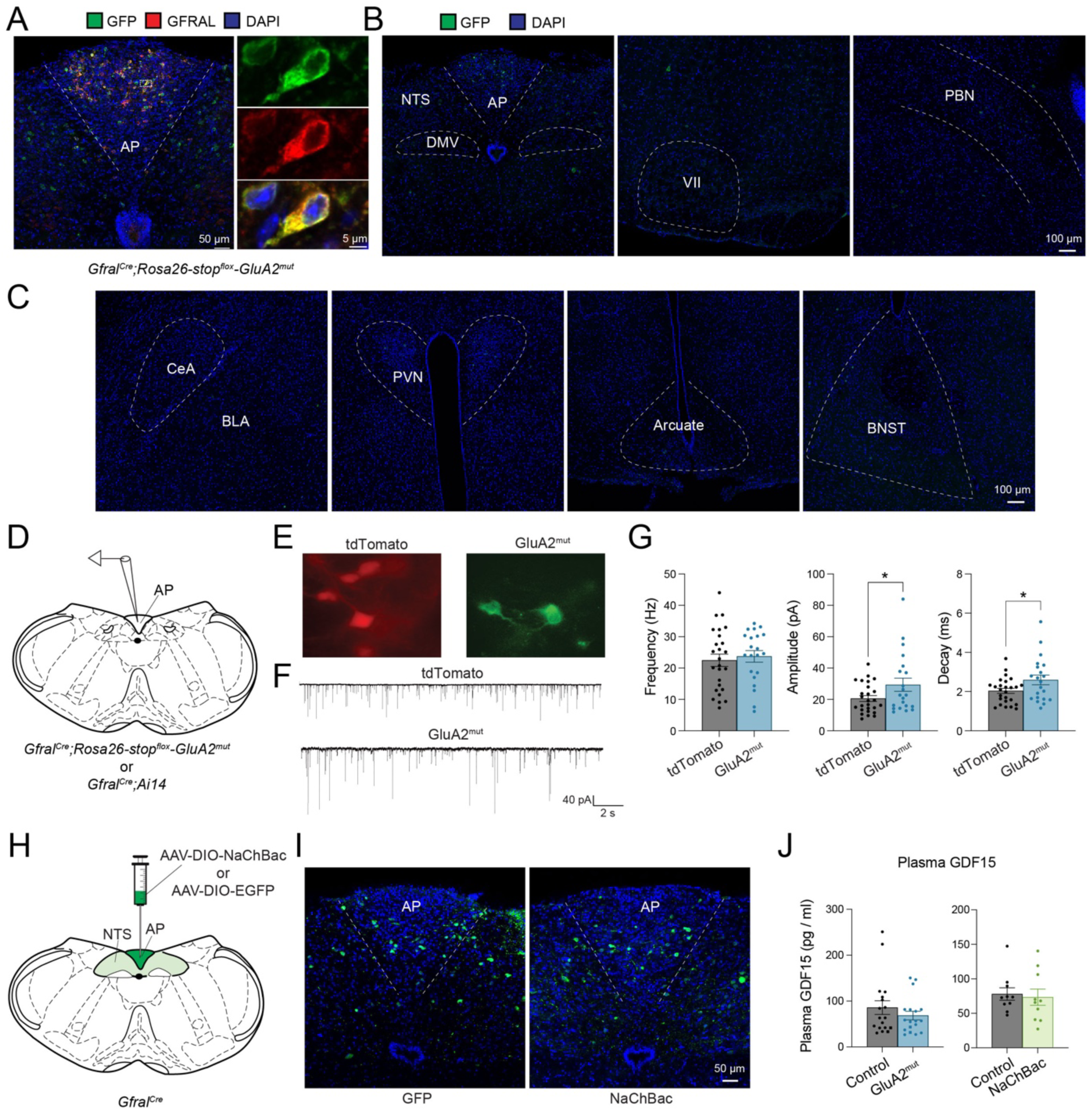
GluA2^mut^ enhanced excitatory transmission in GFRAL neurons. **(A)** Example immunohistochemical images of the AP of GluA2^mut^ mice, showing the expression of eGFP and GFRAL; scale bar, 50 and 5 μm. **(B,C)** Immunohistochemical images of GluA2mut mice, showing the expression of eGFP across different brain regions; scale bar, 100 μm. **(D)** Diagram showing the AP in a coronal brain section for electrophysiological recording. **(E)** Example fluorescence image showing a (left) Gfral-cre:Ai14+ neuron and (right) Gfral-cre:GluA2^mut^+ neuron. **(F)** Representative miniature EPSC traces from GFRAL neurons in control (top) and GluA2^mut^ (bottom) mice. **(G)** Quantification of miniature EPSC frequency (left), amplitude (middle) and decay (right) (control, n = 26 cells / 4 mice, GluA2mut, n = 21 cells / 4 mice; frequency, P = 0.6442 Unpaired t test; amplitude, *P = 0.0450 Mann-Whitney test; decay, *P = 0.0395 Unpaired t test). **(H)** Schematic of NaChBac or eGFP injections in the AP of Gfral-cre mice. **(I)** Example immunohistochemical images of the AP, showing the expression of eGFP or NaChBac; scale bar, 100 μm. **(J)** Plasma GDF15 concentrations in GluA2^mut^ (left) and NaChBac (right) mice (GluA2^mut^: control, n = 18, GluA2^mut^, n = 18; P = 0.4619; Mann-Whitney test) (NaChBac: control, n = 10, NaChBac, n = 10; P = 0.5288; Mann-Whitney test). AP, area postrema; NTS, Nucleus tractus solitarii; PBN, parabrachial nucleus; CeA, central amygdala; BLA, basolateral amygdala; PVN, paraventricular nucleus of hypothalamus; BNST, bed nucleus of the stria terminalis. Data are presented as mean s.e.m.

**Fig. S5.**
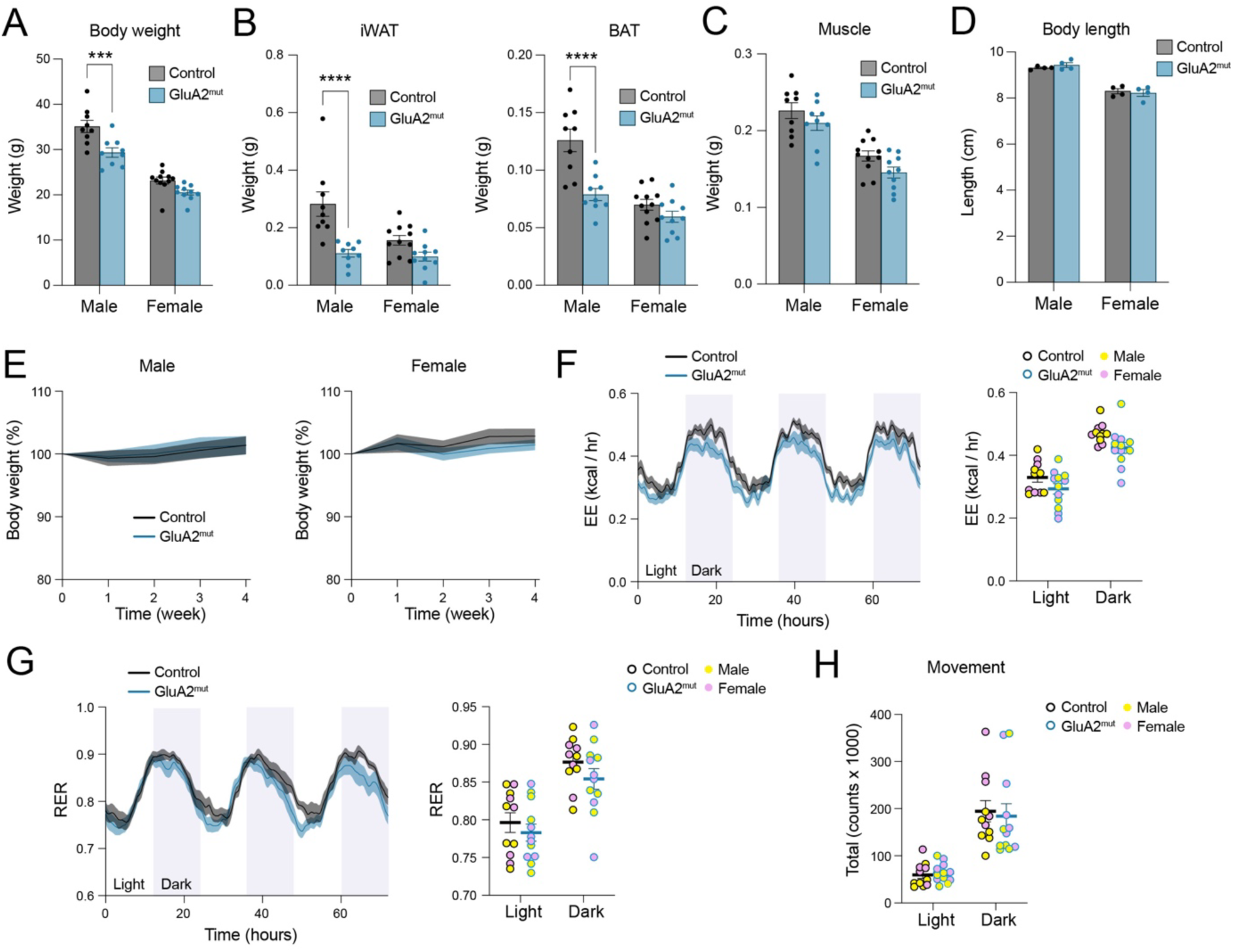
Metabolic effects of GFRAL neuron activation in GluA2^mut^ mice. **(A)** Absolute body weight in male and female mice (male control, n = 9, GluA2^mut^, n = 9; female control, n = 11, GluA2^mut^, n = 10) (male, ***P = 0.0004, female, P = 0.0897; Šídák’s multiple comparisons test). **(B)** Weights of adipose tissue (iWAT male ****P < 0.0001, female P = 0.1730; BAT male ****P < 0.0001, female P = 0.4153; Šídák’s multiple comparisons test). **(C)** Weights of skeletal muscle (male P = 0.3549, female P = 0.1288; Šídák’s multiple comparisons test). **(D)** Body length of male and female mice (male control, n = 4, GluA2^mut^, n = 4; female control, n = 4, GluA2^mut^, n = 4) (male, P = 0.6813, female, P = 0.8609; Šídák’s multiple comparisons test). **(E)** Left, male body weight over time (male; control, n = 8, GluA2^mut^, n = 9; Row factor: F (4, 60) = 2.293 and P=0.0697, Column factor: F (1, 15) = 0.02614 and P=0.8737; two-way repeated-measures (RM) ANOVA followed by Sidak’s post hoc test). Right, female body weight over time (female; control, n = 9, GluA2^mut^, n = 8; Row factor: F (4, 60) = 2.889 and P=0.0297, Column factor: F (1, 15) = 0.7738 and P=0.3929; week 2, *P = 0.0202; two-way repeated-measures (RM) ANOVA followed by Sidak’s post hoc test). **(F)** Energy expenditure (EE) in GluA2^mut^ mice over time (Left) and average EE (Right) (Light, P = 0.1895, Dark, P = 0.0757; Šídák’s multiple comparisons test). **(G)** Respiratory exchange ratio (RER) in GluA2^mut^ mice over time (Left) and average RER (Right) (Light, P = 0.6846, Dark, P = 0.3553; Šídák’s multiple comparisons test). **(H)** Total movement in GluA2^mut^ mice (Light, P = 0.9835, Dark, P = 0.9025; Šídák’s multiple comparisons test). The last 72 hours were analyzed and plotted from the final week in the metabolic cages from GluA2^mut^ mice. Data are presented as mean s.e.m.

**Fig. S6.**
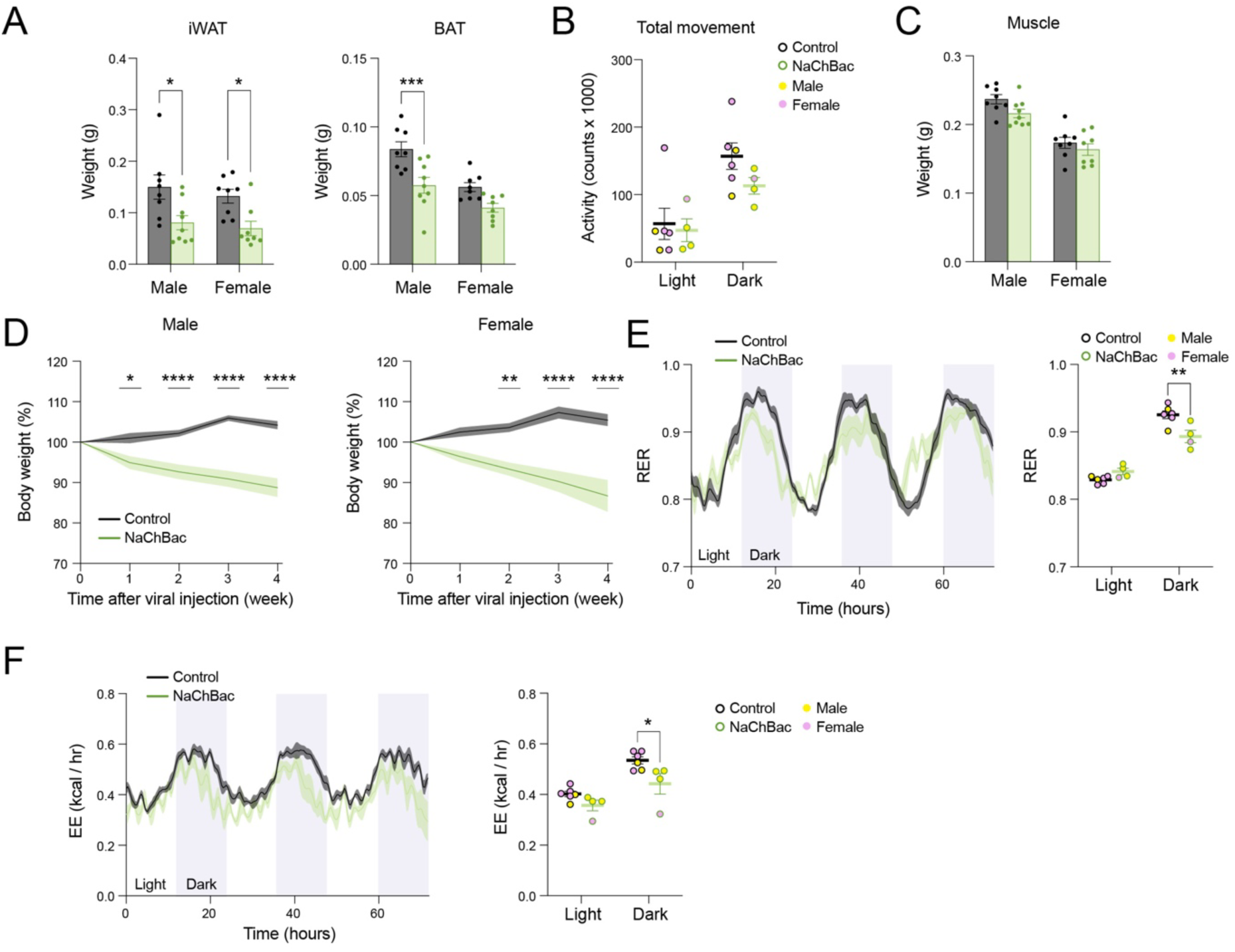
Metabolic effects of GFRAL neuron activation with NaChBac. **(A)** Weights of adipose tissue (male; control, n = 8, NaChBac, n = 9; female; control, n = 8, NaChBac, n = 9) (iWAT male *P = 0.0114, female *P = 0.0267; BAT male ***P = 0.0008, female P = 0.0647; Šídák’s multiple comparisons test). **(B)** Total movement in NaChBac mice (Light, P = 0.9367, Dark, P = 0.2861; Šídák’s multiple comparisons test). **(C)** Weights of skeletal muscle (male P = 0.0994, female P = 0.5864; Šídák’s multiple comparisons test). **(D)** Left, male body weight over time (male; control, n = 8, NaChBac, n = 9; Row factor: F (4, 60) = 3.368 and P=0.0149, Column factor: F (1, 15) = 35.01 and P<0.0001; week 1, *P = 0.0192; week 2, ****P < 0.0001; week 3, ****P < 0.0001; week 4, ****P < 0.0001; two-way repeated-measures (RM) ANOVA followed by Sidak’s post hoc test). Right, female body weight over time (female; control, n = 8, NaChBac, n = 8; Row factor: F (4, 56) = 2.507 and P=0.0522, Column factor: F (1, 14) = 28.84 and P<0.0001; week 1, P = 0.1057; week 2, **P = 0.0008; week 3, ****P < 0.0001; week 4, ****P < 0.0001; two-way repeated-measures (RM) ANOVA followed by Sidak’s post hoc test). **(E)** Respiratory exchange ratio (RER) in NaChBac mice over time (Left) and average RER (Right) (control, n = 6, NaChBac, n = 4; Light, P = 0.2601, Dark, **P = 0.0018; Šídák’s multiple comparisons test). **(F)** Energy expenditure (EE) in NaChBac mice over time (Left) and average EE (Right) (Light, P = 0.2800, Dark, *P = 0.0140; Šídák’s multiple comparisons test). The last 72 hours were analyzed and plotted from the final week in the metabolic cages from NaChBac mice. Data are presented as mean s.e.m.

**Fig. S7.**
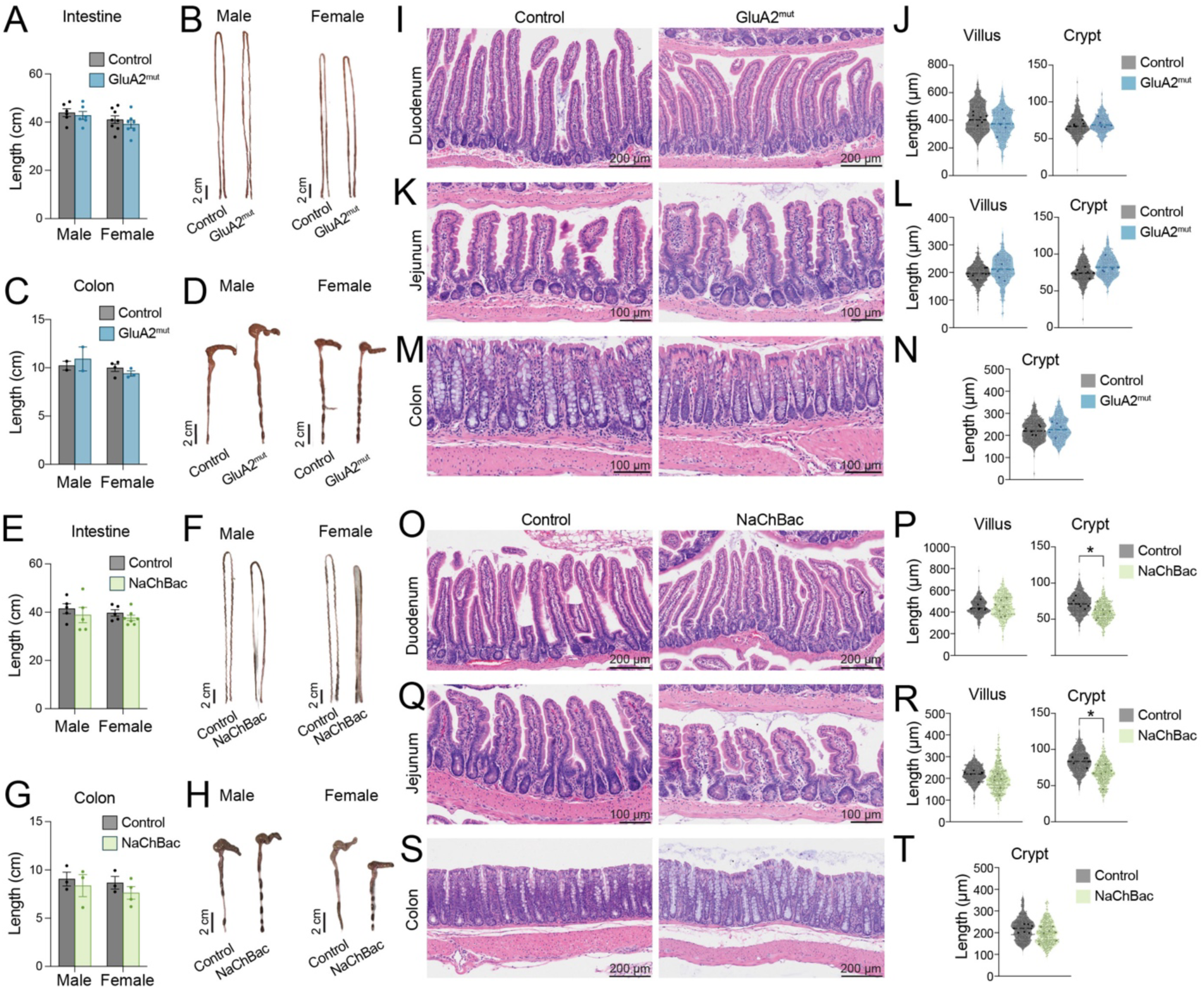
Changes in intestine and colon after GFRAL neuron activation. **(A)** Intestinal length in GluA2^mut^ male and female mice (male control, n = 6, GluA2^mut^, n = 6; female control, n = 8, GluA2^mut^, n = 7) (Male, P = 0.8769, female, P = 0.6443; Šídák’s multiple comparisons test). **(B)** Representative intestine from male and female GluA2^mut^ mice; scale bar, 2 cm. **(C)** Colon length in GluA2^mut^ male and female mice (male control, n = 4, GluA2^mut^, n = 5; female control, n = 4, GluA2^mut^, n = 3) (Male, P = 0.8595, female, P = 0.7638; Šídák’s multiple comparisons test). **(D)** Representative colon from male and female GluA2^mut^ mice; scale bar, 2 cm. **(E)** Intestinal length in NaChBac male and female mice (male control, n = 5, NaChBac, n = 5; female control, n = 5, NaChBac, n = 7) (Male, P = 0.6030, female, P = 0.7002; Šídák’s multiple comparisons test). **(F)** Representative intestine from male and female NaChBac mice; scale bar, 2 cm. **(G)** Colon length in NaChBac male and female mice (male control, n = 3, NaChBac, n = 3; female control, n = 3, NaChBac, n = 4) (Male, P = 0.8198, female, P = 0.5927; Šídák’s multiple comparisons test). **(H)** Representative colon from male and female NaChBac mice; scale bar, 2 cm. **(I)** Representative HE images from duodenum in GluA2^mut^ mice; scale bar, 200 μm. **(J)** Quantification of villus and crypt length from duodenum (control, n = 6, GluA2^mut^, n = 5) (Villus P = 0.2882; Crypt, P = 0.6245; Unpaired t test). **(K)** Representative HE images from jejunum in GluA2^mut^ mice; scale bar, 100 μm. **(L)** Quantification of villus and crypt length from jejunum (Villus P = 0.4464; Unpaired t test; Crypt, P = 0.1255; Mann-Whitney test). **(M)** Representative HE images from colon in GluA2^mut^ mice; scale bar, 100 μm. **(N)** Quantification of crypt length from colon (Crypt, P = 0.6215; Unpaired t test). **(O)** Representative HE images from duodenum in NaChBac mice; scale bar, 200 μm. **(P)** Quantification of villus and crypt length from duodenum (control, n = 6, NaChBac, n = 7) (Villus P = 0.8357; Mann-Whitney test; Crypt, *P = 0.0217; Unpaired t test). **(Q)** Representative HE images from jejunum in NaChBac mice; scale bar, 100 μm. **(R)** Quantification of villus and crypt length from jejunum (Villus P = 0.1807; Mann-Whitney test; Crypt, *P = 0.0101; Unpaired t test). **(S)** Representative HE images from colon in NaChBac mice; scale bar, 100 μm. **(T)** Quantification of crypt length from colon (Crypt, P = 0.1497; Unpaired t test). **(J, L, N, P, R, T)** Each small point represents the length of an individual crypt or villus, and each larger point indicates the average value for an individual mouse. Statistical analyses were performed on the per-mouse averages. Data are presented as mean s.e.m.

**Fig. S8.**
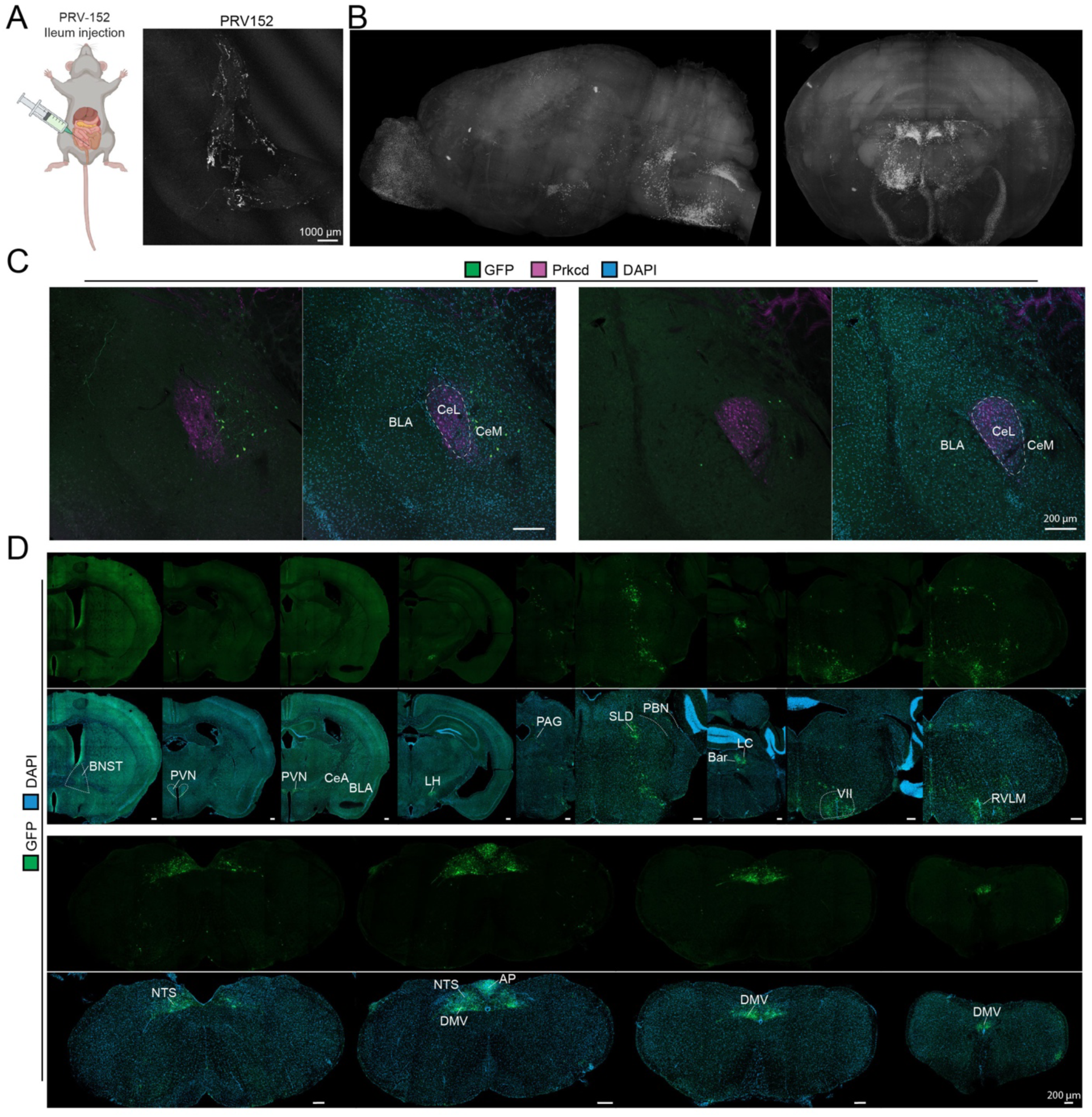
PRV-mediated transsynaptic tracing from the intestine to the brain. **(A)** Left, schematic overview of PRV injections in the small intestine (n = 4). Right, immunohistochemical image of the small intestine with PRV expression; scale bars, 1000 μm. **(B)** Whole mouse brain image of PRV+ cells in the brain 5 days after PRV injection in the small intestine. **(C)** Immunohistochemical images of the CeA 5 days after PRV injections in the small intestine, stained for GFP and Prkcd; scale bars, 200 μm. **(D)** Immunohistochemical images of the brain, 5 days after PRV injections in the small intestine, stained for GFP; scale bars, 200 μm. CeL, central amygdala lateral; CeM, central amygdala medial; BLA, basolateral amygdala; BNST, bed nucleus of the stria terminalis; PVN, paraventricular nucleus of hypothalamus; CeA, central amygdala; LH, lateral hypothalamus; PAG, periaqueductal gray; SLD, sublaterodorsal nucleus; PBN, parabrachial nucleus; Bar, barrington’s nucleus; LC, locus ceruleus; VII, facial motor nucleus; RVLM, rostral ventrolateral medulla; NTS, Nucleus tractus solitarii; AP, area postrema; DMV, dorsal motor nucleus of the vagus. Created in BioRender. van de Lisdonk, D. (2026) https://BioRender.com/dey02x5.

**Fig. S9.**
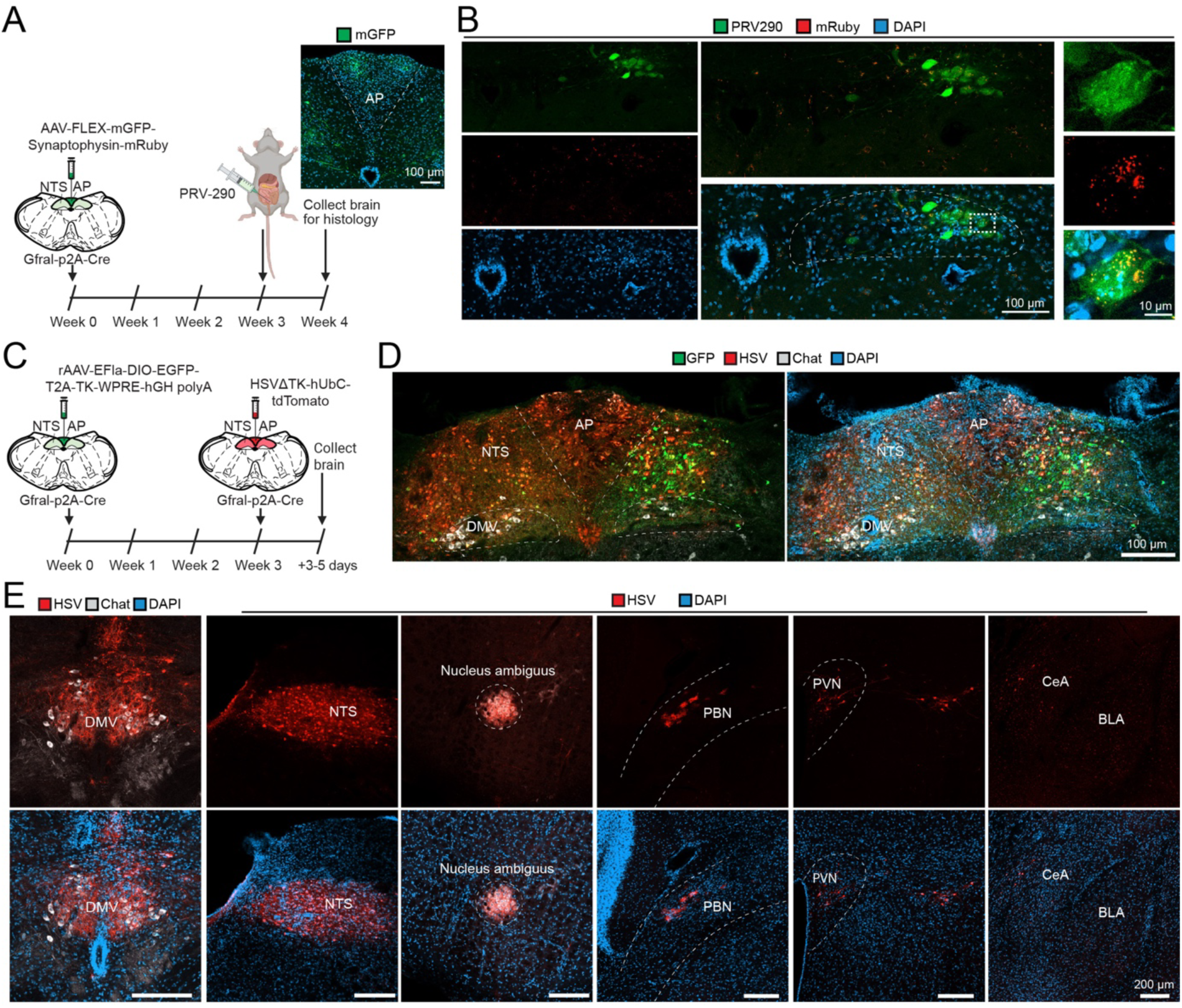
GFRAL neurons project to ChAT⁺ neurons in the dorsal motor nucleus of the vagus. **(A)** Timeline of the experiments, with AAV-Synaptophysin injections on week 0 in Gfral-p2A-Cre mice and PRV injection in the small intestine on week 3. And Immunohistochemical images of the AP after AAV-Synaptophysin-mGFP injections; scale bars, 100 μm. **(B)** Immunohistochemical images of the DMV, 4 weeks after AAV-Synaptophysin in the AP of GFRAL-p2A-Cre mice and 4 days after PRV-290 injections in the small intestine, stained for PRV-290 and mRuby; scale bars, 100 and 10 μm (n = 2). **(C)** Timeline of the experiments, with rAAV-EFla-DIO-EGFP-T2A-TK-WPRE-hGH polyA injections on week 0 in the AP of Gfral-p2A-Cre mice and HSV injection on week 3 (n = 3). **(D)** Immunohistochemical images of injection site 3 days after the final HSV injection, stained for GFP, HSV and Chat; scale bars, 200 μm. **(E)** Immunohistochemical images of the brain, 3-5 days after HSV injections, stained for HSV and Chat; scale bars, 200 μm. AP, area postrema; NTS, Nucleus tractus solitarii; DMV, dorsal motor nucleus of the vagus; PBN, parabrachial nucleus; PVN, paraventricular nucleus of hypothalamus; CeA, central amygdala; BLA, basolateral amygdala. Created in BioRender. van de Lisdonk, D. (2026) https://BioRender.com/dey02x5.

**Fig. S10.**
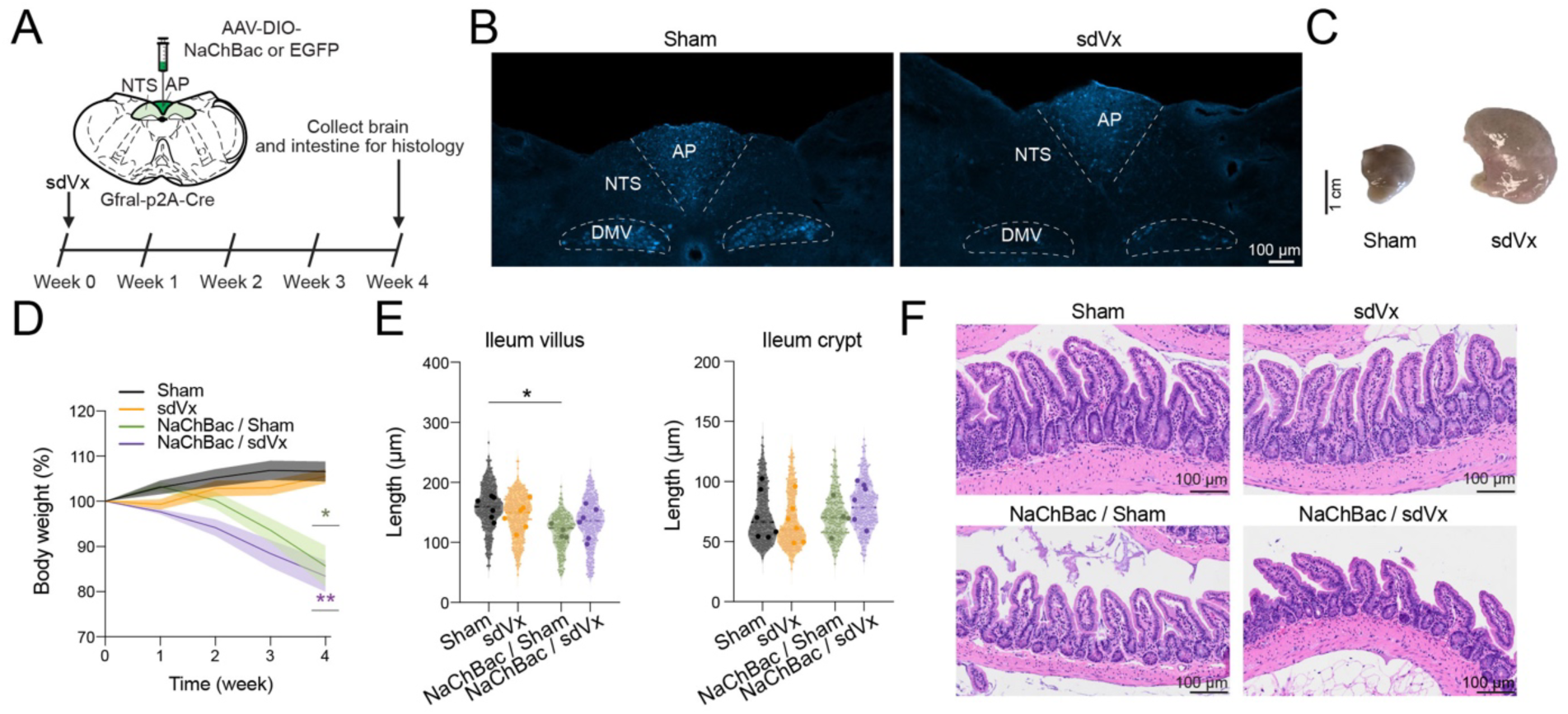
GFRAL neurons regulate intestinal type 2 immune responses via the vagus nerve without affecting body weight or intestinal villus length. **(A)** Schematic overview of sdVx and NaChBac injections into the AP of GFRAL-p2A-Cre mice. **(B)** Staining of fluorogold in the DMV sham (left) and vagotomised (right) mice to confirm efficiency of subdiaphragmatic vagotomy (sdVx). **(C)** Anatomical verification of sdVx, showing normal (left) and enlarged stomachs (right). **(D)** Body weight over time in sham (n = 6) vs sdVx (n = 6) vs NaChBac+sham (n = 4) vs NaChBac+sdVx (n = 6) mice (Row factor: F (1.626, 29.27) = 11.40 and P=0.0005, Column factor: F (3, 18) = 12.67 and P=0.0001; week 4 Sham vs. sdVx, P = 0.9751; Sham vs. NaChBac + Sham, *P = 0.0338; Sham vs. NaChBac + sdVx, **P = 0.0012; Two-way repeated-measures (RM) ANOVA followed by Sidak’s post hoc test). **(E)** Quantification of villus (left) and crypt (right) length in sham vs sdVx vs NaChbac+sham vs NaChBac+sdVx. Villus; (Sham vs. sdVx; P = 0.6590, Sham vs. NaChBac + sham; *P = 0.0435, Sham vs. NaChBac + sdVx; P = 0.1337; Šídák’s multiple comparisons test). Crypt; (Sham vs. sdVx; P = 0.9346, Sham vs. NaChBac + sham; P = 0.9599, Sham vs. NaChBac + sdVx; P = 0.6224; Šídák’s multiple comparisons test). Each small point represents the length of an individual crypt or villus, and each larger point indicates the average value for an individual mouse. Statistical analyses were performed on the per-mouse averages. **(F)** Representative HE images from Ileum in mice; scale bar, 100 μm. Data are presented as mean s.e.m.

**Fig. S11.**
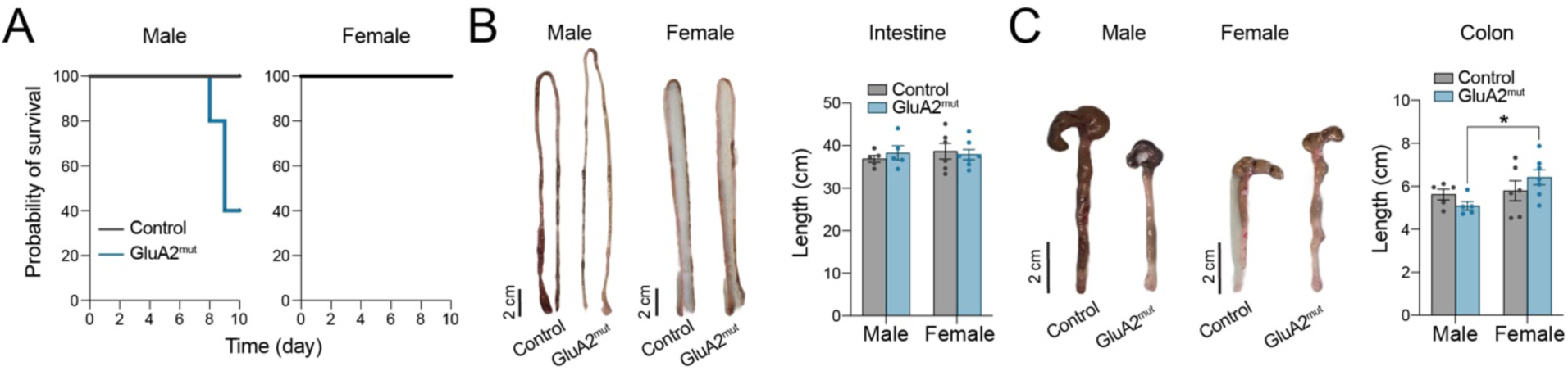
GFRAL neuron activation differentially affects body weight in male and female mice during DSS-induced colitis. **(A)** Survival curves of control and GluA2^mut^ male **(left)** and female **(right)** mice after 2% DSS treatment (Male: P = 0.0511; Mantel-Cox test) (Female: P > 0.9999; Mantel-Cox test). **(B)** Intestinal length in male and female GluA2^mut^ mice after 2% DSS treatment (male, P = 0.7740, female, P = 0.9078; Šídák’s multiple comparisons test). **(C)** Colon length in male and female GluA2^mut^ mice after 2% DSS treatment (male vs female GluA2^mut^, *P = 0.0304; Šídák’s multiple comparisons test). Data are presented as mean s.e.m.

